# Persistence of tobacco-mutated alveolar progenitor cells after smoking cessation mirrors long term risk of lung adenocarcinoma

**DOI:** 10.64898/2026.07.06.736766

**Authors:** Moritz J. Przybilla, Amany Ammar, Hugh Selway-Clarke, Andrew R. J. Lawson, Michael Spencer Chapman, Hyunchul Jung, Kate H. C. Gowers, Pantelis A. Nicola, Marie-Belle El Mdawar, Manuela Platé, Kate E. J. Otter, Zoe C Hagel, Chuen R. Khaw, Inigo Martincorena, Adam Pennycuick, Peter J. Campbell, Sam M. Janes

## Abstract

Tobacco smoke shapes mutations, selection and clonal expansion in lung epithelial cells. Smoking cessation leads to divergent epidemiology in the two most common lung cancers: squamous cell carcinoma risk declines sharply, while adenocarcinoma risk is preserved. To investigate this discrepancy, we analysed 806 genomes of alveolar type II (AT2) cells and found persistently elevated mutation burdens after cessation. In contrast, in the proximal airway, rare basal stem cells with near-normal mutation burden expand after cessation, protecting against squamous cell carcinoma. Targeted single-molecule DNA sequencing of AT2 cells revealed positive selection for *TP53* and cell cycle and MAPK genes, supporting continued cancer risk. A multistage carcinogenesis model emphasised the importance of a small population of hypermutated cells in the alveoli and reproduced the divergent epidemiological trajectories following cessation due to distinct regenerative dynamics. Our findings suggest that differences in mutational burden and clonal regeneration explain post-cessation trends in lung cancer subtypes.

**One sentence summary:** Cancer risk reflects not only cumulative exposure to toxins, but the capacity of tissues to erase genomic damage through regeneration from protected cells with clonal advantage.

## Introduction

Somatic mutations occur daily in trillions of cells in human tissues through endogenous processes or environmental insults. Most are inconsequential, but occasional mutations provide a competitive advantage to a cell, leading to clonal expansion (*1*). In sun-exposed skin, for example, up to 25% of cells carry ultraviolet-induced, cancer-driving mutations causing expanding mutant clones (*2*). Some of these mutations are known to cause cancer, although a single cell likely needs to acquire multiple mutations before transformation. Distinct selective pressures, including lifestyle or environmental factors, may accelerate the competitive expansion of clones over time promoting the acquisition of additional advantageous alterations and driving a somatic evolutionary process.

Lung cancer exemplifies this evolutionary model, with tobacco exposure as a dominant mutagen (*3*). As the leading cause of cancer mortality worldwide, understanding its development is critical. Non-small cell lung cancer (NSCLC) is the major subtype, and this largely comprises of squamous cell carcinoma (LUSC), arising proximally in airway basal cells, and adenocarcinoma (LUAD), originating from alveolar type 2 (AT2) cells in the lung periphery (*4–6*).

While both subtypes are associated with smoking, the last 30 years has seen a change in their relative incidence with adenocarcinoma growing as a proportion (*4*, *7*). Smoking rates have declined in this period, and there has been a growing number of ex-smokers (*8*, *9*). Notably, these two lung cancer subtypes differ markedly in their epidemiological trajectories following smoking cessation: the risk of squamous cell cancer of the airways declines (*9–12*), whilst adenocarcinoma risk remains elevated. This is an unexplained observation (*4*).

In the past, we and others have demonstrated that normal basal cells in the airways of cigarette smokers are dominated by tobacco-related mutations (*13*, *14*). After smoking cessation, highly mutated airway basal cells are progressively replaced by the progeny of a small population of basal cells with no or low tobacco-related mutation burden, possibly recruited from a protected progenitor niche (*15–17*). To date, the genomic damage accumulating in human alveoli cells and the accompanying clonal dynamics have not been characterised.

Here, we address whether the maintained risk of LUAD compared to falling risk of LUSC following smoking cessation is explained by the persistence of highly mutated AT2 cells in the lung parenchyma as opposed to the progressive replacement of high mutant basal cells by rare low mutant cells in the airways. We integrate whole-genome and single-molecule sequencing of normal alveolar cells with classical multistage models of carcinogenesis, testing whether differences in mutation burden and post-cessation repopulation dynamics could account for this epidemiological observation.

## Results

### Smoking and non-small cell lung cancer subtype frequency

The link between smoking and lung cancer is well established. We mapped the distinct epidemiological trajectories of NSCLC subtypes following smoking cessation within the Surveillance, Epidemiology and End Results (SEER) registry data (*18*) to confirm previously documented changes in NSCLC subtype rates (*12*), with squamous cell carcinoma falling since 2000 with the rate of adenocarcinoma progressively increasing despite falling current smoking rates. The proportion of non-neuroendocrine NSCLCs that were adenocarcinoma rose from 28% in 1988 to 64% in 2020 (**Fig. 1A**).

**Fig. 1.**
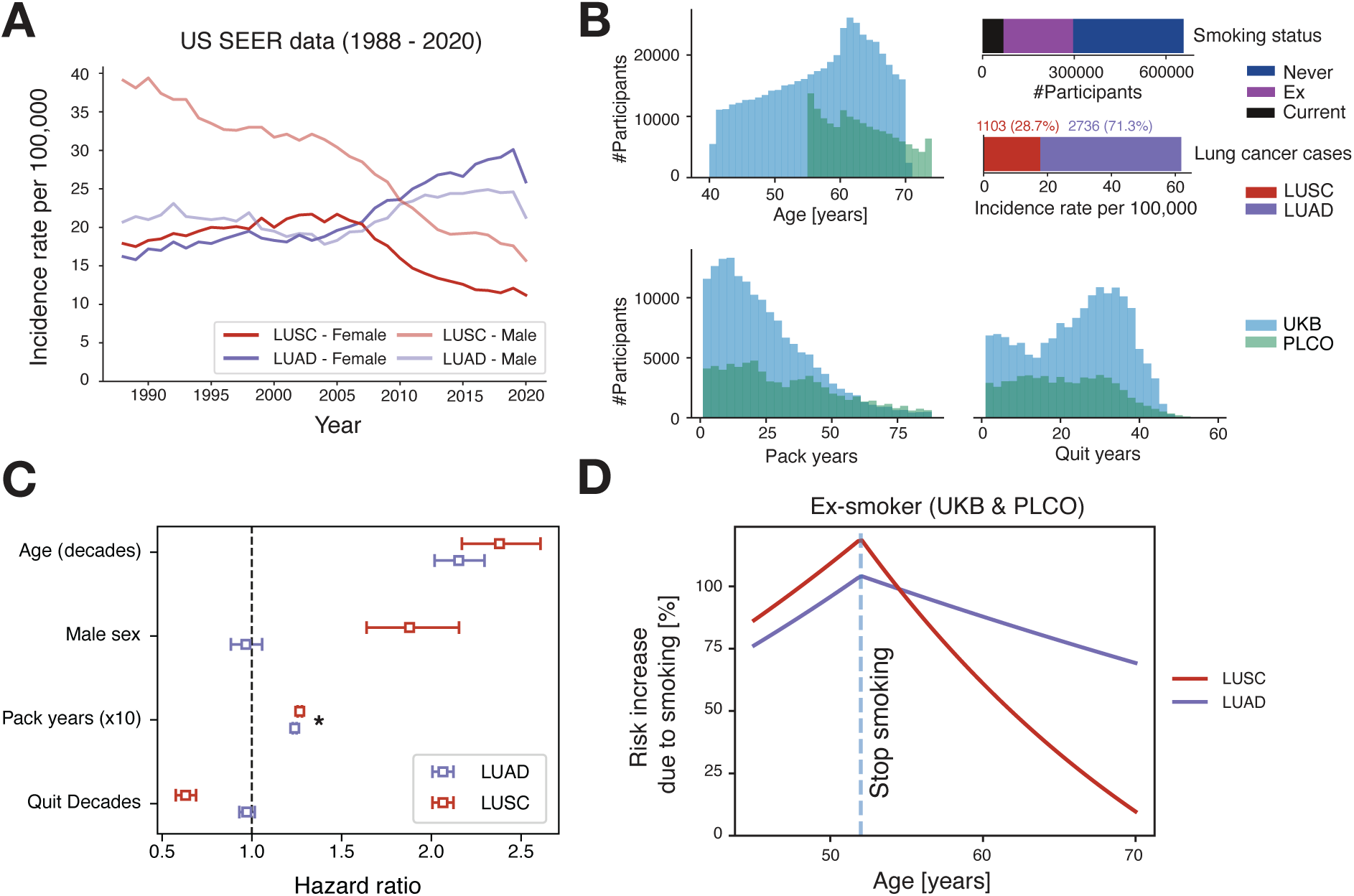
The intricate relationship between tobacco smoking and lung cancer risk. (**A**) Age-adjusted rates of Lung Adenocarcinoma (LUAD) and Lung Squamous Cell Carcinoma (LUSC) in the US SEER 8 dataset between 1988 and 2020, stratified by sex. (**B**) Cohort overview of age, smoking status, lung cancer cases as well as pack and quit year distributions in UKB and PLCO. (**C**) Effects of incrementing age, male sex, pack years and years since cessation of smoking (in years and pack years) on the risk (hazard) of getting each form of cancer, fit by a Cox proportional hazard model. The asterisk indicates small confidence intervals being present for pack-years. (**D**) Illustration of proportional increase in hazard due compared to a never smoker for each histology over time, as fit in the Cox model, for a woman who quits smoking at 52 years (the 90^th^ percentile of quit ages).

We examined epidemiological data from ~650,000 individuals in the UK Biobank and Prostate, Lung, Colorectal, Ovarian (PLCO) cancer screening trial cohorts to further investigate temporal trends of squamous and adenocarcinoma incidence (*19*, *20*). The combined cohorts had 2,736 LUAD (71.3%) and 1,103 LUSC (28.7%) cases, with wide variation in the number of pack-years of smoking and years since smoking cessation in ex-smokers across the cohort (**Fig. 1B**, **fig. S1A,B**). We applied a multivariate survival model to evaluate the effects of smoking cessation on subtype risk. As expected, smoking and age contributed to increased risk for both LUAD and LUSC (**Fig. 1C**, **fig. S1C,D**). Strikingly, smoking cessation decreased subsequent risk of LUSC by a third per decade of cessation (HR: 0.68, 0.61-0.74 95% CI) but had minimal effect on risk of lung adenocarcinoma (HR: 0.96, 0.92-0.99, 95% CI).

We can illustrate the magnitude of this difference by estimating the risk of adenocarcinoma versus squamous cell carcinoma over time for a hypothetical female who starts smoking at 18 and stops smoking at age 52: by age 70, her model-predicted risk of LUSC has almost returned to approximately that of a non-smoker, but her risk of adenocarcinoma remains 75% higher (**Fig 1D**). These data suggest that smoking cessation has fewer benefits in reducing risk of adenocarcinoma compared to squamous cell carcinoma.

### Mutation burden of alveolar type 2 cells

How the divergence in cancer type risk is related to biology in the lung is unclear. To investigate, we characterised the mutational landscape of normal AT2 cells and compared it to our previously reported data from bronchial basal epithelial cells (*13*). We established single-cell derived alveolar organoids from 9 individuals who had undergone lung resections for cancer (3 never-, 3 ex- and 3 current-smokers; **Fig. 2A**, **fig. S2, table S1**). In brief, we dissected and enzymatically digested lung parenchyma biopsies to generate single-cell suspensions, from which AT2 cells were isolated (*21*). We developed a protocol to expand single AT2 cells into clonal organoids, which then underwent WGS (**Methods**). Immunofluorescence, whole-mount staining, lysotracker staining, transmission electron microscopy and gene expression confirmed organoids were derived from AT2 cells (**Fig. 2B**, **fig. S3**). Colony-forming efficiency ranged from 5-10% and was lower in older patients and ex-smokers (p<0.05 and p<0.01, respectively, Wilcoxon-rank sum test; **fig. S3H-I**). We performed WGS across 806 organoids (57-96 organoids/patient; **table S2**). 97% (785/806) of organoids had a monoclonal origin by their variant allele fraction (VAF) distribution and were retained for downstream analysis (**fig. S4A**). Several sister organoids (from the same clonal expansion) were sequenced from one patient as a control, showing minimal accumulation of mutations during *in vitro* culture (~50 mutations; **fig. S4B**). Data from the alveolar organoids were compared to single cell-derived colonies of basal cells from the proximal airways of 16 patients with varying smoking history, analysed using the same pipeline (*13*).

**Fig. 2.**
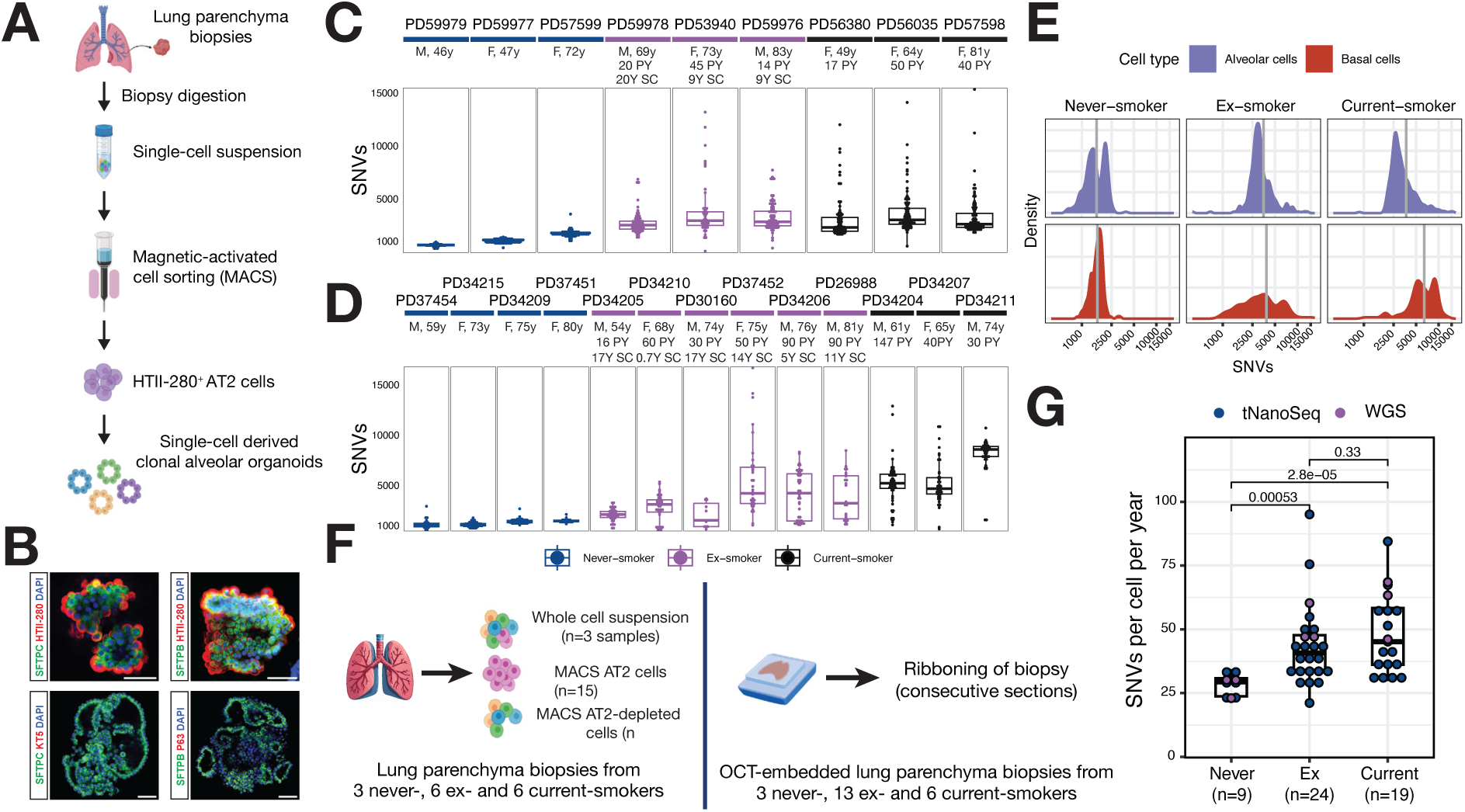
The mutation burden in stem cells in the proximal airway and alveoli of the lung. (**A**) Schematic of the experimental process underlying the generation of clonal alveolar organoids. A full schematic is provided in fig. S2. (**B**) Whole-mount staining of clonal alveolar organoids for AT2 cell markers (SFTPC, SFTPB and HTII-280) and basal cell markers (TP63, KRT5 and ABCA3). The organoids did not show expression of basal cell markers TP63 and KRT5. Scale bar = 50 µm. (**C**) (top) Box plot depicting the total number of single nucleotide variants (SNVs) per AT2 cell (organoid) split according to the respective patient. Patients are ordered according to smoking status and age. Sex, age and smoking cessation as well as pack year history is annotated if applicable. (**D**), Equivalent figure to (**C**) showing the re-analysis from basal cell colonies from Yoshida et al., 2020. (**E**), Density plot contrasting the mutation burden of AT2 (top) and basal cells (bottom) across smoking status. The mean mutation burden across each category is annotated as vertical grey line. (**F**), Experimental overview of samples used for tNanoSeq. For the 15 lung parenchyma biopsies, 3 whole cell suspensions, 15 MACS AT2 cell suspensions and 3 MACS AT2-low cell suspensions were sequenced. (**G**) Number of SNVs per genome per year from tNanoSeq and whole genome sequencing (WGS) for all samples included in the study. Dots are coloured according to sequencing type, with smoking categories being depicted individually. The number of samples per category is indicated below. Annotated p-values represent the results of a Wilcoxon-rank sum test between the indicated groups (SC: smoking cessation; PY: pack-year).

AT2 cells of ex- and current-smokers had a significantly higher overall burden of somatic mutations (51 and 60 SNVs/cell/year respectively) than never-smokers (28 SNVs/cell/year; p<0.01, Wilcoxon-rank sum test; **Fig. 2C-D**). In both ex- and current-smokers, the distribution of SNV burden was highly skewed, with a long right tail indicative of a small subpopulation (~5–15%) of cells with markedly elevated burdens exceeding 5,000 SNVs per cell (**Fig. 2E**). This contrasted with the narrower and more symmetric distribution observed in never-smokers. Similar right-skewed patterns were observed for dinucleotide variants (DNVs) and indels, with significantly elevated burdens in ex- and current-smokers (p<0.01; **fig. S5A**). These findings suggest that tobacco exposure heterogeneously affects alveolar cells, with a subset bearing a disproportionate burden.

A key observation from our previous study of proximal airways was that, in ex-smokers, 20-40% of basal epithelial cells had near-normal mutation burdens and no mutational signatures of tobacco exposure. This population of cells was highly expanded compared to the 2-4% of such cells in current smokers (*13*). This was not the case in the alveolar organoids – there was insufficient evidence, as tested using a pairwise per patient Kolmogorov-Smirnov, that the distributions of mutation burdens between current smokers and ex-smokers differed substantially, despite the ex-smokers having quit 9-20 years prior (p>0.05 across patient-pairs, KS test; **Fig. 2C-E**, **fig. S5B-D; table S1**).

To validate these findings in an extended cohort, we applied targeted nanorate sequencing (tNanoSeq), a newly developed high-fidelity single-molecule sequencing method (*22*), to measure the burden of SNVs and indels in AT2 cells from 37 individuals (**Fig. 2F–G**, **fig. S5E**). Unlike clonal organoid sequencing, tNanoSeq provides a population-agnostic readout, reflecting the average mutational burden across all cells in the sample. Consistent with organoid data, we observed no significant difference in average mutation burden between ex- and current-smokers (p=0.33, Wilcoxon rank-sum test), but markedly lower burdens in never-smokers (p=5.3×10^−4^ and p=2.8×10^−5^, for ex- and current-smokers, respectively).

Leveraging the combined cohort, we found a significant dose–response relationship between smoking exposure and mutation burden in alveolar cells, as measured by linear regression (p<0.001; **fig. S5F**). For every 10 pack-years of smoking, the average mutation burden increased by ~11%. Given that a normal alveolar cell in a never-smoker accumulates approximately 28 mutations per year, this corresponds to ~1,400 mutations by age 50. A 10 pack-year exposure - equivalent to 3,650 cigarette packs - would thus contribute an additional ~150 mutations per cell, or roughly one new mutation in every AT2 cell per 24 packs smoked. Notably, due to the pronounced right skew of the distribution, some alveolar cells likely acquire mutations at much higher rates, approaching one mutation per cigarette pack in extreme cases (**Fig. 2E**).

Of note, we observed structural variants (SVs) in ~25% of AT2 organoids (**table S3**; **fig. S6**). Most SVs were deletions (*23*) in known fragile sites in the genome, including *PTPRD*, *LRP1B* or *FHIT* (**fig. S6C**). In keeping with other normal tissues, we found few copy number alterations (CNAs), predominantly focal losses (**fig. S7**).

### Mutational signatures of alveolar cells

Mutational signatures reflect distinct patterns of DNA mutations which inform the underlying biological processes or exposures responsible for their generation. To define the lasting genomic imprint of smoking and distinguish it from endogenous damage, we extracted mutational signatures from the combined dataset of the alveolar and the proximal airway epithelial cells published previously (*13*). The most prevalent mutational processes were the clock-like mutational processes SBS5 and SBS40 (**Fig. 3**). Clock-like signatures accumulate steadily over time - reflecting ongoing endogenous processes such as cell division or DNA repair – and are ubiquitously present across both healthy and malignant tissues (*24*, *25*).

**Fig. 3.**
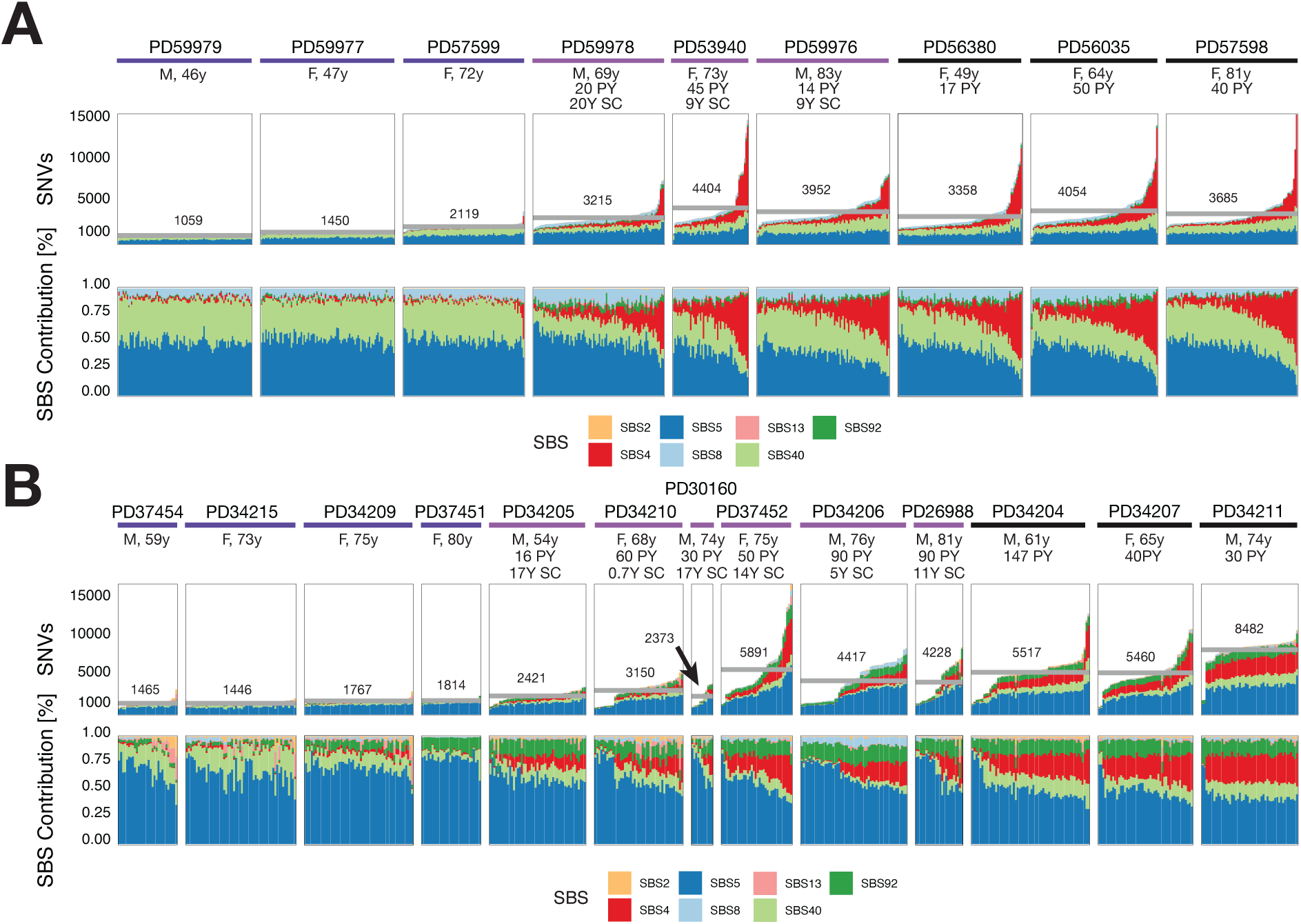
The mutational signatures present in alveolar stem cells in relation to smoking. (**A**) (top) Bar plot depicting the total number of single nucleotide variants (SNVs) per AT2 cell as highlighted coloured by single-base pair substitution (SBS) signature contribution as well as the relative contribution of SBS signatures (bottom). Bar plot is split according to the respective patient, with cells being ordered according to increasing mutation burden. Patients are ordered according to smoking status and age. The horizontal grey line indicates the mean burden calculated across all organoids of a patient. Sex, age and smoking cessation as well as pack year history is annotated if applicable. The arrow indicates the mean burden for PD30160. (**B**), Equivalent figure to (**A**) showing the re-analysis from basal cell colonies from Yoshida et al., 2020. SC: smoking cessation; PY: pack-year.

Among current and ex-smokers, the excess SNV burdens in AT2 cells compared to never-smokers were predominantly driven by COSMIC signature SBS4, the major signature associated with tobacco mutagenesis (*24–30*). This observation was generally supported at the level of DNVs and indels, associated with tobacco-associated signatures DB2 and ID3 (**fig. S8**). Within a given current or ex-smoker, the considerable cell-to-cell variance in mutation burden observed was almost entirely accounted for by variable SBS4 burdens, with some cells carrying 100-200 SBS4 mutations and others carrying many thousands (**Fig. 3B**). In contrast to the proximal airways, where ex-smokers had a significant population of basal cells with near-normal mutation burden and absence of SBS4, there was no appreciable difference in the prevalence or contributions of SBS4 to AT2 cells between current and ex-smokers.

Signatures associated with APOBEC mutagenesis, SBS2 and SBS13, are found in >50% of adenocarcinomas (*24*), but were noticeably absent from normal alveolar cells. This suggests APOBEC is a cancer-specific mutational process acting late in the decades-long process of carcinogenesis, similar to observations in the oesophagus (*31*, *32*). SBS8, previously linked to late replication errors (*33*), accounted for 5-10% of mutations in alveolar cells. SBS92 has been previously linked to tobacco smoke exposure in healthy bladder urothelium and bladder cancer (*28*, *34*) and was seen in both smokers and never-smokers in proximal airway basal epithelium. However, it was absent from AT2 cells despite accounting for 5-20% of cells in proximal airways.

### Genomic imprints of the cell of origin in genomes of AT2 cells

Positive selection for non-synonymous coding mutations is widespread across normal tissues, especially those cell types that maintain lifelong proliferative potential (*35*). To assess the strength of selection acting on AT2 cells, we estimated the dN/dS ratio, which measures the excess or depletion of non-synonymous mutations compared to expectation, corrected for mutational and genomic context (*36*). In our WGS data from AT2 organoids, we estimated the exome-wide dN/dS ratio to be 1.02 (CI_95%_=0.98-1.05) – with a ratio of 1.0 denoting neutrality, this estimate suggests that selection is not a strong force shaping the clonal architecture of alveolar cells. In contrast, for proximal airway organoids, the estimated exome-wide dN/dS ratio was 1.08 (CI_95%_=1.04-1.12; **Fig. 4A**), consistent with 1 in 13 non-synonymous mutations being under positive selection in bronchial epithelium.

**Fig. 4.**
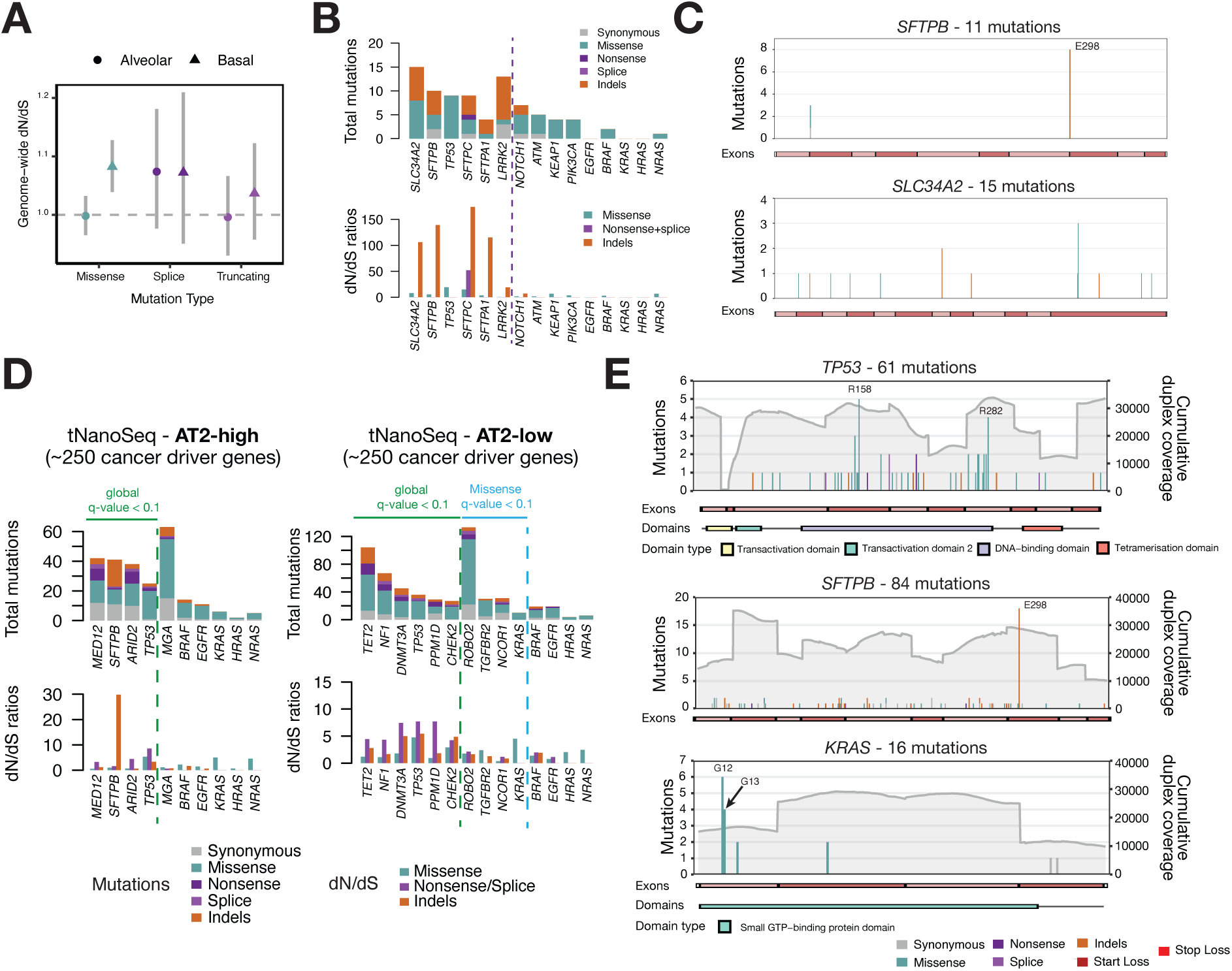
Selective landscape and genomic imprints of LUAD cell of origin in alveolar stem cells. **(A)** Genome-wide dN/dS ratio for missense, nonsense, splice and truncating mutations in alveolar and basal cells from WGS. The horizontal dotted line highlights a dN/dS ratio of 1. 95% confidence intervals for each dN/dS estimate are highlighted by grey vertical lines. **(B)** (top), Number and consequence of mutations detected in AT2 organoids. Mutations are coloured according to impact, including synonymous, missense, nonsense and splice mutations as well as indels (bottom), Observed-to-expected ratios for missense substitutions, truncating (nonsense and essential splice site) substitutions, and indels. Genes left of the vertical dotted line are under significant positive selection (dndscv, q<0.1), whereas genes right of the line are known LUAD drivers. **(C)** Gene representation for *SFTPB* and *SLC34A2* for mutations found across single cell-derived alveolar type 2 (AT2) organoids. Number of mutations are annotated on the y-axis. Exon annotations are shown on the x-axis. (**D**) Equivalent visualisation as shown in (**B**) for samples dominated by AT2-high (left) and AT2-low (right) cells from tNanoSeq. **(E)** Gene representation equivalent to (**C**) for *TP53*, *SFTPB* and *KRAS* across all samples analysed with tNanoSeq. In addition, the cumulative duplex coverage per site is shown as a grey line and according to the scale on the right y-axis. Exon as well as protein annotation are shown if available for the respective transcript.

At the level of individual genes, we identified 5 that had a significant excess of mutations in AT2 cells after correction for multiple hypothesis testing (q<0.1; **Fig. 4B**). Of these, only *TP53* is a known cancer gene – showing the expected pattern of missense mutations clustered in the DNA-binding domain (q=0.0026; **fig S9A**).

Two of the other significantly mutated genes encode surfactant proteins: *SFTPB* and *SFTPC* (q=9.2×10^−5^, 9.1×10^−6^, respectively). The other surfactant protein gene *SFT1A1* was also highly mutated, though not significant after multiple hypothesis testing correction (q=0.2). Interestingly, the same surfactant genes are frequently mutated in lung adenocarcinomas (*37*, *38*). These mutations are thought to arise from a mutational process enriched for indels within the genomic footprint of highly expressed genes, potentially resulting from transcription–replication conflicts (*37–39*). Consistent with this model, we observe a sequence-context–driven indel hotspot in *SFTPB* (E298), indicating that this genomic region exhibits an elevated mutation rate that is further amplified by high transcriptional activity. These data suggest that adenocarcinomas deriving from these cells may inherit pre-existing patterns of mutational clustering in surfactant genes. A similar phenomenon is seen in the albumin gene, *ALB*, in both hepatocellular carcinomas (*38*) and normal liver cells (*40*, *41*).

We found similar patterns of indel accumulation in *SLC34A2* and *LRRK2* (q=5.2×10^−8^ and 2.1×10^−2^ respectively; **Fig. 4B-C**, **fig. S9B-C**). Both *SLC34A2*, a gene implicated in surfactant recycling (*42*) and *LRRK2* are highly expressed in AT2 cells (*43–45*). An increase of indels in the introns of those genes (**fig. S9D**) as well as high dN/dS estimates for indels specifically, suggest these results to possibly represent lineage-defining mutational imprints.

### Mutations in known cancer genes in alveolar cells

Lung adenocarcinomas have frequent mutations activating key signalling pathway genes, including *KRAS* and *EGFR* (33% and 14% respectively) (*4*). Previous studies have reported driver mutations in these genes to be widespread in normal alveolar tissue (*46*), but our genome-wide data from 806 AT2 organoids identified only 2 mutations in *BRAF*, 1 in *NRAS* (**Fig. 4B**) and none in *KRAS* or *EGFR*. Only 1 of the *BRAF* mutations represented a canonical hotspot (D594G). This provides a sharp contrast to the driver-rich mutation landscape of organoids grown from the squamous epithelium of proximal airways (**fig. S9E**).

To increase statistical power to detect positive selection in alveolar cells, we used tNanoSeq of ~250 cancer genes (**Fig. 2F**) (*22*). Each dx (duplex coverage) thereby reflects the equivalent of 0.5 single cell-derived organoids. We profiled genomic DNA from magnetic-activated cell sorted AT2 cells from a total of 15 patients, including the 9 patients used to create clonal organoids. To complement this, we also sequenced unsorted single cell suspensions before (n = 3); the AT2-low cell fraction after sorting (n = 3); and frozen parenchymal biopsies from 22 patients with diverse smoking histories. For analysis, we combine these into an ‘AT2-low’ sample group, given that they all contain limited AT2 cells in comparison to the majority of AT1 and parenchyma infiltrating immune cells, including alveolar macrophages and T cells (*47–49*) (**fig. S10**; **table S1**).

We detected 22,070 mutations across all samples, with a mean duplex coverage of 664 dx (range = 40-2,204 dx; **table S4**). AT2-high samples demonstrated statistically significant excess of mutations in *SFTPB* and *TP53*, in keeping with the whole genome data, and *MED12* and *ARID2* also under positive selection (**Fig. 4D**). There was no obvious association with smoking status. Indels for *SFTPB* were located at the identical position (E298) compared to the single cell data, underscoring the mechanism of microhomology-mediated indel formation which generate clustered frameshift events. Mutations in the mediator complex subunit 12 (*MED12*) are frequent in breast fibroadenomas and uterine leiomyosarcomas, where hotspot mutations in codons L36 and G44 disrupt the interaction between MED12 and CDK8 (*50*). However, in AT2-sorted cells, we instead observed excess nonsense or indel mutations across the gene. *ARID2,* involved in chromatin remodelling, is part of the SWI/SNF complex and commonly inactivated in lung adenocarcinoma (*51*, *52*). In AT2-low samples (**Fig. 4D**), we found 6 positively selected genes: *TET2*, *NF1*, *DNMT3A*, *TP53*, *CHEK2* and *PPM1D* (q<0.1). Interestingly, all these genes are drivers of clonal haematopoiesis of indeterminate potential (*53*). We believe that this signal results from a combination of blood cells in the pulmonary circulation and alveolar macrophages derived from blood monocytes.

Examining individual hotspots in the genome, we found 16 mutations in *KRAS*, of which 10 were in G12 or G13, the most frequently mutated residues in lung adenocarcinoma (**Fig. 4F**). However, it should be noted that these mutations occurred in very small fractions of cells in each sample and were only detected in ever-smokers (<<1% cells; **fig. S10B-C**). Several other individual sites in the genome showed positive selection, including known hotspots in *PIK3CA*, *SF3B1*, *BRAF*, *EP300* and *FGFR2* (*54*, *55*) (**Supplementary Data**).

Taken together, our data suggest that driver mutations are infrequent in normal AT2 cells, accounting for <1% cells, even in individuals who smoke tobacco. This contrasts with the proximal airway epithelium, where 10-25% of cells carry drivers in older individuals with a smoking history (*13*), and also other normal tissues such as skin, blood, endometrium and oesophagus (*2*, *31*, *56–58*). Nonetheless, in keeping with multistage models of carcinogenesis, occasional alveolar cells do carry rare driver mutations, and these include some of the canonical variants seen in adenocarcinomas.

### Regenerative potential across lung compartments explains subtype incidence rates post smoking cessation

To evaluate whether the divergent epidemiological trajectories of squamous cell carcinoma and adenocarcinoma in ex-smokers could be explained by differences in regenerative potential between their respective cells-of-origin, we developed a simulation framework grounded in multistage carcinogenesis. Specifically, we hypothesised that the rapid decline in LUSC risk after smoking cessation - contrasted with the persistent LUAD risk - could be driven by repopulation of airway basal cells with near-normal mutation burden, a process not observed in alveolar cells.

We modelled cancer risk as the probability of a single cell acquiring *k* rate-limiting driver mutations over time, with driver mutation rates (*r*) derived from observed empirical distributions (*59*). In this framework, cancer risk increases approximately with *rᵏ*, making the system highly sensitive to differences in mutation burden (**Fig. 5A**). Modest changes in mutation rate *r* can have substantial effects on risk: for *k* = 5, a two-fold increase in rate implies ~32-fold higher risk (*27*). This illustrates why rare high-mutation-rate cells can disproportionately drive tumorigenesis and why this effect is pronounced with increasing *k* (**Fig. 5B**; **figure S11A**). Conversely, even modest effective reductions in *r* - such as those expected from clonal repopulation with near-normal cells - could cause a steep decline in risk. The absence of such repopulation observed in alveolar cells would, in this model, lead elevated risk to persist after smoking cessation, consistent with LUAD epidemiology.

**Fig. 5.**
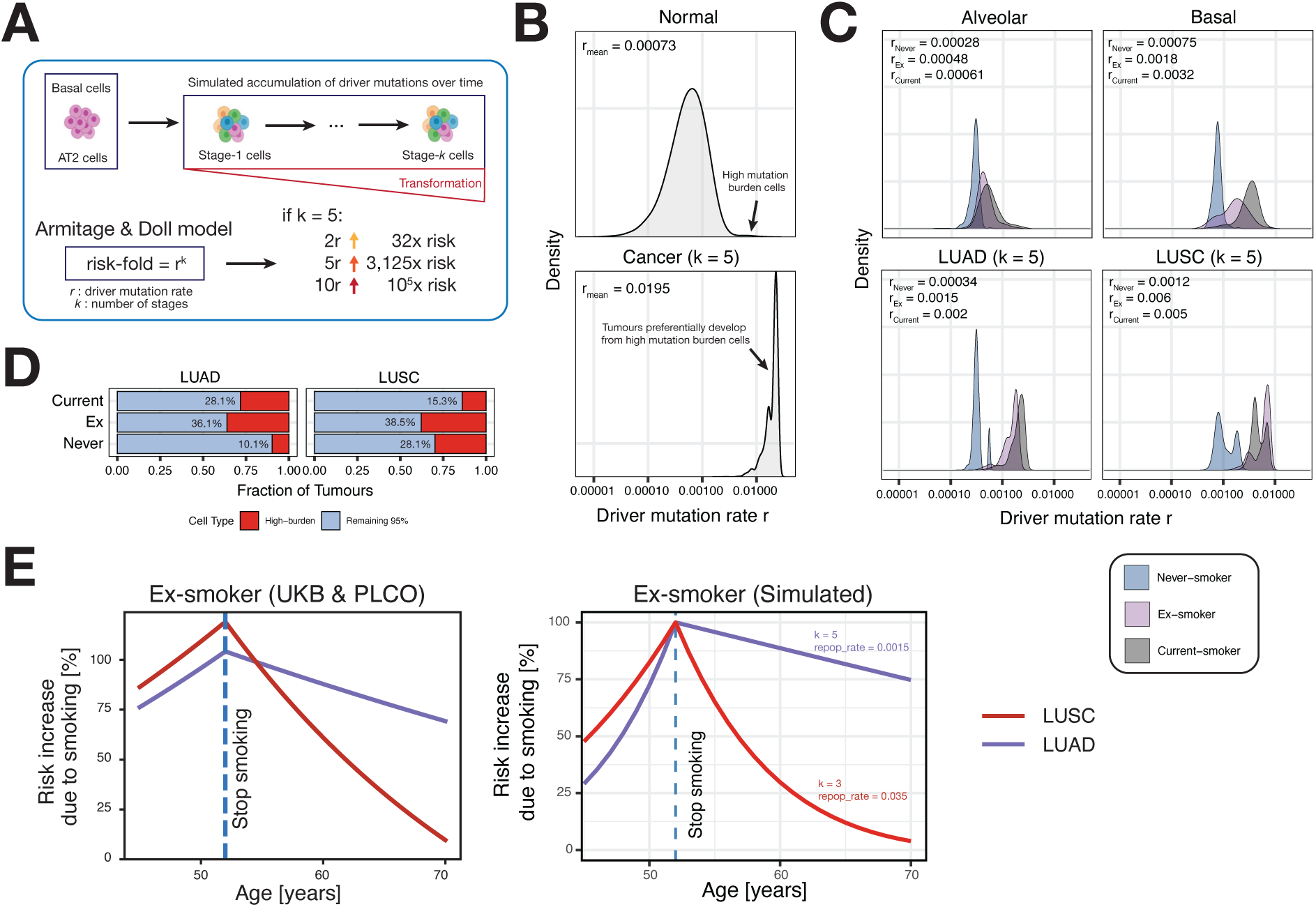
Simulating lung carcinogenesis from genomic damage. (**A**) Schematic principle of the simulation and Armitage-Doll multistage model. (**B**) Simulated driver mutation rate distribution of normal cells (top) and for cancer (bottom) using a gamma distribution with *k* = 5 events. Note a small population of cells with high mutation burden in the top panel. (**C**) (top), Estimated mutation rate *r* distribution for alveolar and basal cells across distinct smoking statuses from real data. (bottom), Simulated tumour origin mutation rate *r* from the empirical data represented in top panel across smoking histories. The visualisation mirrors the conceptually simulated data shown in B. (**D**) Bar plot highlighting the relative risk within each smoking category and cancer type, resulting from 5% high burden cells compared to 95% of the remaining cells. Annotated percentages show the percentage of tumours resulting from hypermutator cells. (**E**) (Left) Visualisation of increase in LUSC and LUAD risk over time as predicted from the Cox Hazard model shown in Fig. 1D. (Right) Simulated risk over time for LUAD (*k* = 5, *repop_rate* = 0.0015) and LUSC (*k* = 3, *repop_rate* = 0.035) according to an Armitage-Doll model. Risk increase is defined as excess subtype-specific risk over a never-smoker.

We applied this model (assuming *k* = 5) to our observed distribution of driver mutation rates (**Fig. 2E**) and we found that the highest mutation-burden cells (defined as the 95^th^ percentile of each smoking specific, empirical distribution) disproportionately drive LUAD and LUSC risk (**Fig. 5C,D**; **fig. S11B,C**). Across smoking categories and cancer subtypes, just 5% of cells accounted for 10.1% to 38.5% of simulated risk. Cell-to-cell variation, as observed in compartments, has therefore a greater effect on total risk compared to the population mean. Due to the contribution of tobacco, mutation rates in normal lung epithelial cells are highly heterogenous; our findings here illustrate how rare, highly mutated clones - though a minority - can dominate cancer initiation.

Finally, we extended this framework to simulate across-lifespan cancer risk trajectories for ex-smokers via a post-cessation gradual replacement of the stem cell population by cells drawn from never smoker mutational burden distributions (**Fig. 5E**; **fig. S11D**; Methods). Possible values for the rate-limiting driver mutation count *k* and repopulation rates were informed by previous studies (*27*, *36*, *60*, *61*). We found that the observed decline in LUSC risk after cessation could be qualitatively recapitulated by an annual repopulation rate of ~3.5% yr^−1^ and *k* = 3 rate-limiting driver events (**Fig. 5E**). This rate was consistent with the replacement rates observed by Yoshida and colleagues (median = ~3.5% yr^−1^, range = 0.4 – 33.7%) (*13*). In contrast, the modest reduction in LUAD risk was consistent with a ~23-fold lower rate (0.15% yr^−1^), aligning with a lack of regeneration in alveolar tissue (**Fig. 5E**; **fig. S11D**). Together, these subtype-specific differences in mutation rate distributions and regenerative dynamics provide a plausible mechanistic explanation for the divergent epidemiology of lung cancer subtypes following smoking cessation.

## Discussion

Smoking cessation is the most effective intervention to reduce lung cancer risk (*4*, *8*, *62*). The benefits are almost immediate: lung squamous cell carcinoma incidence declines steeply in the years after quitting. Yet, in stark contrast, lung adenocarcinoma risk remains substantially elevated for decades in ex-smokers. This divergence, documented across epidemiological cohorts, presents a biological paradox: the somatic mutations caused by tobacco smoke are irreversible and heritable — so why does risk fall in one compartment of the lung but persist in the other?

Our results offer clonal dynamics as a parsimonious explanation for this epidemiological paradox. By comparing the mutational landscapes of normal bronchial basal cells and AT2 cells, we reveal striking differences in how these epithelial compartments respond to tobacco exposure and cessation. In basal cells, smoking generates highly mutated basal cell clones, but some progenitor cells which carry substantially lower mutation loads are also present and expand following cessation. Alveolar tissue shows no evidence of this clonal reset. There is no low mutant population of AT2 cells in smokers or ex-smokers and mutational damage remains high even in long-term ex-smokers, with a subset of cells carrying an extremely elevated mutation burden. Importantly, our model does not require invoking protected niches, differential DNA repair, or ongoing mutagenesis after cessation; the observed epidemiological trends emerge naturally from differences in tissue dynamics and clonal competition alone (*63*). Extrapolating from these findings, we propose a conceptual model in which the benefit of smoking cessation depends on whether damaged epithelial compartments can reset their clonal composition. Tissues with high regenerative turnover and cell replacement can rapidly reduce cancer risk if they have a residual population of cells with limited numbers of mutations, whereas tissues lacking this capacity retain a long-term “mutational memory” of past exposures. This framework reconciles the irreversible nature of somatic mutation with the reversible component of cancer risk observed epidemiologically.

Phenotypically normal clones must typically acquire multiple driver events before any single cell can initiate malignancy (*1*, *64*). The selective landscapes of alveolar and basal cells support this model. Basal cells exhibit strong positive selection across numerous genes, including established LUSC drivers, reflecting a regenerative environment that tolerates the accumulation of oncogenic mutations. In contrast, alveolar cells show sparse selection beyond *TP53*, with signals largely confined to cell cycle and MAPK pathways. This limited selective pressure aligns with their persistence post-cessation - additional driver mutations cannot be afforded and could destabilise the compartment. Within this framework, a lower number of rate-limiting steps (*k*) in LUSC compared to LUAD would match the observed biology.

More broadly, this work highlights the value of integrating normal-tissue mutational data with evolutionary theory to explain population-level cancer trends. By linking tissue-specific clonal dynamics to subtype-specific incidence patterns, our study provides a mechanistic framework for understanding how carcinogen exposure, tissue regeneration, and somatic evolution interact to shape long-term cancer risk.

Finally, our results underscore the foundational role of somatic mutation and selection in lung carcinogenesis. Tobacco exposure imposes a powerful selective pressure on epithelial clones, whose expansion and persistence differ across compartments. These dynamics, rooted in tissue architecture and stem cell biology, create distinct evolutionary trajectories within the same organ - illustrated by the divergent behaviour of basal and alveolar lineages in LUSC and LUAD. A fuller understanding of cancer risk will require large-scale efforts to quantify clonal growth and competition in normal tissues over time, linking mutational burden to lineage fate and disease emergence.

## Acknowledgments

We thank the patients who consented to the use of their tissue for clinical research. We thank CASM Support (L. O’Neill, C. Hardy, R. Thatcher), CASM Research Managers (I. Tang, H. Robertson, M. Venables), and CASM IT (A. Butler, A. Menzies, A. Burdett, V. Offord) for their contributions. We are grateful to the CASM Support Laboratory (K. Roberts, T. Baxter, K. Smith, E. Ferla, M. Khan, L. Allen) for assistance with sample handling and shipments. We thank H. Savin and Y. Hooks for histopathology and laboratory support throughout the project.

This work was supported by a Cancer Research UK programme grant. S.M.J. is supported by Cancer Research UK programme grants (EDDCPGM\100002 and EDDPGM-Nov25/100001) and an MRC Programme grant (MR/W025051/1). S.M.J. also receives support from the CRUK Lung Cancer Centre and the CRUK City of London Centre Award (C7893/A26233), as well as the CRUK ACED Award to UCL. Additional support was provided by the Rosetrees Trust, the Garfield Weston Trust, and the University College London Hospitals Charitable Foundation. I.M. was supported by Cancer Research UK (C57387/A21777), the Milky Way Research Foundation, the Dr Josef Steiner Cancer Research Foundation, and the Wellcome Trust. This work was partly undertaken at UCLH/UCL, which receives funding from the Department of Health’s NIHR Biomedical Research Centre scheme. MP was supported by The Rosetrees Trust, Asthma and Lung UK, and the National Institute for Health Research University College London Hospitals Biomedical Research Centre. Several researchers on this grant were funded in whole, or in part, by the Wellcome Trust [Grant number 220540/Z/20/A]. For the purpose of Open Access, the author has applied a CC BY public copyright licence to any Author Accepted Manuscript version arising from this submission.’

## Supplementary Materials

### Materials and Methods

#### Experimental methods

##### Human lung tissue collection

Human samples were sourced ethically from University College London Hospitals following ethical approval from the National Research Ethical Committee (REC reference 18/SC/0514) (IRAS no. 245471). Written informed consent was obtained from all donors. Lung parenchyma was taken from surgical samples from patients undergoing cancer resections, at least 3 cm from the tumour, following macroscopical assessment by an experienced pathologist. All samples were used in accordance with the informed consent.

##### Human alveolar organoid culture

Human feeder-free alveolar organoids were generated and maintained as previously described (*1*). Briefly, pleura and any visible airways were removed before dissecting human lung parenchymal samples into small pieces of ~1 mm each. Dissected tissues were incubated with collagenase type I (GIBCO, 17100-017), dispase (Corning, 354235) and DNaseI (Thermo Fisher Scientific, 10104159001) for 60 min at 37 °C. Digested mixtures were filtered through 70 µm and 40 µm cell strainers, followed by lysis of red blood cells with ACK lysis buffer (Thermo Fisher Scientific, A1049201). The cells were washed with DMEM/F12 (Thermo Fisher Scientific, 12634028) containing 10% FBS and resuspended in primary HTII-280 antibody (Terrace Biotech, TB-27AHT2-280) diluted 1:50 in FACS buffer (1 mM EDTA and 1% FBS in PBS). The mixture was incubated at 4 °C in the dark for 60 min. The cells were then washed twice with FACS buffer and incubated with anti-Mouse IgM MicroBeads (Miltenyi, 130-047-301) diluted 1:5 in FACS buffer for 30 min at 4 °C in the dark. The cells were washed twice with FACS buffer, resuspended in an appropriate volume of FACS buffer and loaded onto pre-prepared Miltenyi LS columns (Miltenyi, 130-042-401) to be collected magnetically. MACS-sorted AT2 cells were counted and resuspended in serum free feeder free organoid medium described below and then mixed with equal volumes of Matrigel. 50 μl drops of cells-medium-Matrigel mixture, at a density of 5000 cells/drop, were plated in each well of a 24-well plate. Drops were allowed to solidify at 37 °C for 1 hr. Subsequently, 500 μl of medium was added to each well. The medium is composed of advanced DMEM/F12 supplemented with 10µM SB431542 (Abcam, Ab120163), 3µM CHIR99021 (Tocris, 4423), 1µM BIRB796 (Tocris, 5989), 1X ITS (Thermo Fisher Scientific, 41400045), 15mM HEPES (Thermo Fisher Scientific, 15630080), 1X GlutaMAX (Thermo Fisher Scientific, 35050061), 5µg/ml Heparin (Sigma-Aldrich, H3149), 1X antibiotic antimycotic solution (Sigma-Aldrich, A5955), 50ng/ml recombinant human EGF (Thermo Fisher Scientific, PHG0313), 1X B27 (Thermo Fisher Scientific, 17504044), 1X N2 (Thermo Fisher Scientific, 17502048), 1X N-acetylcystein (Sigma-Aldrich, A9165), 10µM Y27632 (Cambridge Bioscience, Y1000), 10ng/ml recombinant human FGF10 (BioLegend, 559304), recombinant human BMP4 10ng/ml (BioLegend, 595202). Media were changed every 3 days and organoids were grown for 21 days. Organoids with an area greater than 500 μm^2^ were counted at day 15 of culture to determine colony forming efficiency (CFE). Organoids were collected after 21 days of culture. Cultures were regularly tested for mycoplasma.

##### Generation of clonal alveolar organoids, DNA extraction and library preparation

Clonal alveolar organoids were generated using the limiting dilution method. AT2 cells were seeded at low cell densities of 20, 50, 100, and 200 cells per medium/Matrigel drop. After growth for 21 days, single organoids were expanded for 2 passages to create clonal cultures as follows. Individual organoids were manually picked with the help of a stereomicroscope, transferred into fresh drops in separate wells of 12-well plates, and dissected with needles into 5-6 pieces (*2*). Drops were allowed to solidify at 37 °C for 1 hr before being covered with medium. After growth for 21 days, grown organoids were further passaged by mechanical dissociation using narrowed glass pipettes, replated in a 1:2-1:4 ratio, and allowed to expand for 21 days (**fig. S2**). To enhance the reproducibility of sequencing purely clonal organoids, only a single organoid from each established clonal culture was utilised for WGS (**fig. S2**). Genomic DNA was extracted from organoids according to manufacturer’s instructions using the Arcturus PicoPure DNA extraction kit (Thermo Fisher Scientific, KIT0103). Briefly, a single organoid from each clonal culture was transferred into a separate well of a skirted 96-well PCR plate (Eppendorf, 0030129512) prefilled with 20 μl proteinase K solution, then incubated at 65 °C for 6 hr. Proteinase K was inactivated at 75 °C for 30 min. The plates were stored at −20 °C until library preparation. Libraries were prepared for low-input DNA sequencing as previously described (3) and only organoids with a DNA concentration ≥ 5 ng/μl were retained for sequencing.

##### Cytospin preparations

To validate the identity and purity of the MACS-sorted AT2 cells, cytospins were prepared from the whole cell suspension before MACS, MACS-sorted AT2, and flowthrough. After cell counting, cells were centrifuged at 400 g for 5 min and resuspended in PBS at a cell concentration of 7.5×10^5^ cells per 1 ml. 150,000 cells (200 μl) were loaded into an assembled cytospin cassette and spun at 800 rpm for 5 min with high acceleration in the cytospin machine. Cassettes were removed carefully from the machine, and the slides were left to air-dry for 20 min. The slides were incubated in 2% PFA for 10 min at room temperature and washed twice with PBS. The samples were then immunostained with antibodies against specific AT2 cell markers, HTII-280 and SFTPC, using the same protocol employed to immunostain alveolar organoids as described below.

##### Immunofluorescence staining

Organoids were first incubated with organoid harvesting solution (R&D, 3700-100-01) for 1 hr at 4°C to dissolve Matrigel and then fixed with 4% paraformaldehyde (PFA) for 30 min on ice. Organoids were cryo-embedded in optimal cutting temperature compound (OCT, CellPath KMA-0100-00A) and stored at −80°C before cryosectioning at 8 μm thickness. For antigen retrieval, cryosections were immersed in 10 mM sodium citrate buffer pH 6.0 and allowed to boil for 5 min. Slides were then washed twice with PBS before being permeabilized with 0.3% Triton X-100 in PBS for 10 min. Slides were incubated with blocking solution (5% normal donkey serum (NDS), 1% bovine serum albumin (BSA), and 0.1% Triton X-100 in PBS) for 1 hr at room temperature, followed by incubation with primary antibodies (**table S5**) at 4 °C overnight. Slides were washed three times with PBS and then incubated with secondary antibodies (**table S6**) for 2 hr at room temperature. After three washes with PBS, samples were mounted in Vectashield with DAPI (Vector labs). Stained organoids were imaged using a Leica DMi8 microscope and images were analysed using ImageJ v2.1.0.

##### Whole mount staining

Whole mount staining of clonal organoid cultures was carried out as previously described (*4*). Organoids were fixed with 4% PFA for 30 min on ice, washed twice with PBS, resuspended in permeabilization solution (0.3% Triton X-100 in PBS), and then transferred to a low attachment 24-well plate. Organoids were incubated for 20 min at room temperature, followed by incubation with blocking solution (5% NDS, 1% BSA and 0.1% Triton X-100 in PBS) for 1 hr at room temperature. Organoids were then incubated with primary antibodies (**table S5**) for 48 hr at 4 °C. After washing three times with washing buffer (0.1% Triton X-100 and 0.5% BSA in PBS), organoids were incubated with secondary antibodies (**table S6**) overnight at 4 °C. After three washes, organoids were stained with DAPI for 15 min. Organoids were washed twice and incubated with a fructose-glycerol clearing buffer overnight at 4 °C before being mounted on a microscope slide. Slides were imaged using a Leica SP8 and images were analysed using ImageJ v2.1.0.

##### Lysotracker staining

To further characterise alveolar organoids, lysotracker staining was used to stain acidic components of lamellar bodies. Lysotracker (Invitrogen, L12492) was diluted in pre-warmed SFFF medium to a final concentration of 50 ng/μl and then organoid drops were incubated in situ with the diluted Lysotracker solution for 30 min at 37 °C. Lysotracker solution was then removed, and organoid drops were carefully washed with PBS for 5 min. Fresh SFFF medium was added and images were captured using an EVOS live imaging microscope (*5*).

##### Electron microscopy

Organoids were fixed with 2% paraformaldehyde/1.5% glutaraldehyde prepared in 0.1 M sodium cacodylate buffer pH 7.4 for 30 min at room temperature. After washing twice with 0.1 M sodium cacodylate buffer, organoids were osmicated with 1% osmium tetroxide and 1.5% potassium ferricyanide for 1 hr at 4°C and then washed multiple times with 0.1 M sodium cacodylate buffer. Organoids were post-fixed with 1% tannic acid in 0.05 M sodium cacodylate buffer for 45 min at room temperature and then rinsed with deionised water before being dehydrated in graded ethanol (70%/90%/100%/100% dry) twice in each for 5 min at room temperature. Samples were embedded in Epon resin, allowed to polymerize overnight at 60 °C and sectioned using an ultramicrotome (Leica UC7). 70 nm thick sections were collected onto formvar-coated slot grids. Grids were stained with Reynolds lead citrate, rinsed with deionised water, and left to air dry. Images were acquired using a Tecnai T12 Spirit Biotwin (Thermofisher, USA) equipped with Morada CCD camera and iTEM software (EMSIS GmbH, Germany) (*6*).

##### RNA isolation and qRT-PCR

Alveolar organoids were first retrieved from Matrigel, and then total RNA was extracted according to the manufacturer’s instructions using the RNeasy Mini Kit (Qiagen, 74104). Nanodrop 1000 spectrophotometer was used to determine RNA concentration and purity. 0.5 μg of RNA was reverse transcribed using 5X qScript cDNA Supermix (Quantabio, 733-1177) following the manufacturer’s protocol. Quantitative RT-PCR was carried out using TaqMan Fast Advanced Mix (Life technologies, 4444556) and the following TaqMan gene expression assays from Life Technologies: SFTPC (Hs00161628_m1), SFTPB (Hs010900667_m1), SCL34A2 (Hs00197519_m1), and GAPDH (Hs03929097_g1). The mRNA levels of all target genes were quantified using the ΔCt method.

##### DNA extraction from cell suspensions

Genomic DNA was extracted from the spared MACS-sorted AT2 cells of the 9 patients from whom clonal organoids were created, as well as of 6 additional patients (3 current- and 3 ex-smokers). Genomic DNA was also isolated from the unsorted cell suspension before MACS and the AT2-depleted cell suspension from a subset of 3 of those additional patients. DNA was extracted using Qiagen QIAamp DNA Micro Kit (Qiagen, 56304) according to the manufacturer’s instructions and utilised for tNanoSeq.

##### Ribbons preparation and DNA extraction

Ribbons were prepared from 22 OCT frozen parenchymal biopsies from patients with diverse smoking histories and utilised for tNanoSeq. Genomic DNA was isolated in equivalence to the single-cell suspensions using the Qiagen QIAamp DNA Micro Kit (Qiagen, 56304) according to the manufacturer’s instructions and utilised for tNanoSeq.

#### Computational methods

##### DNA sequence alignment

All WGS data were aligned to the GRCh38 reference genome by the Burrows–Wheeler algorithm (BWA-MEM). Targeted Nanorate sequencing was aligned to GRCh37 instead (*7*).

##### Single- & double base-substitution calling

Single nucleotide variants were called using the Cancer Variants through Expectation Maximization (CaVEMan) algorithm (*8*) with copy-number options of major copy number 5, minor copy number 2 and normal contamination 0.1. In cases where samples had a CNA, the ASCAT results were incorporated in the variant calling. In addition to the default ‘PASS’ filter, variants with a median alignment score (ASMD) < 120 and those with a clipping index (CLPM) > 0 were used to remove mapping artefacts. Subsequently, for every mutation identified in any sample from each patient, the number of mutant and wild-type reads were counted using vafCorrect (https://github.com/cancerit/vafCorrect). Further filters described below were applied to identify true somatic mutations and separate them from either germline variants or recurrent sequencing errors.

##### Removing germline variants (binomial filter)

To filter out remaining germline variants, a binomial distribution was fitted to the total variant counts and total depth at each SNV site across all samples from one patient. Thereby, the total depth at the position was utilised as the number of trials with the total number of variant counts as the number of successes. Germline and somatic variants were differentiated based on a one-sided exact binomial test, with the null hypothesis that the number of reads which support the variants across copy number normal samples is drawn from a binomial distribution where p=0.5 (p=0.95 for a copy number equal to one). In contrast, the alternative hypothesis posits that the reads are drawn from a distribution with p<0.5 (or p<0.95). Resulting p-values were corrected for multiple hypothesis testing utilising the Benjamini–Hochberg method and a cut-off was set at q<10−5 to minimise false positives as. Variants for which the null hypothesis could be rejected were classified as somatic, while all others were classified as germline.

##### Removing errors (beta-binomial filter)

Following this, filtering of remaining artefacts was performed by fitting a beta-binomial distribution to the variant counts and depths of all SNVs across samples from the same patient. In principle, the beta-binomial was used as it captures the difference between artefactual variant sites and true somatic variants. Thereby, artefacts often appear to be randomly distributed across samples and can be modelled as drawn from a binomial distribution. True somatic variants will be present at a high VAF in some samples, but absent in others, and are hence best captured by a highly overdispersed beta-binomial. For all variants, the overdispersion parameter (rho) was quantified, with variants that had rho smaller than 0.1 being filtered out as previously described elsewhere (*9*, *10*).

##### Clonality of samples

To estimate the clonal structure within a sample, a truncated binomial mixture model as described previously was applied (*10*). The truncated distribution is used to reflect the minimum number of supporting reads (n = 4) that is required by CaVEMan. In theory, the model will try to separate the overall SNVs into the clones that they could have arisen from, each with their own probability (VAF) and proportion (the amount of variants a clone contributes). The proportion of cells that inhabit a clone can be approximated by twice the estimated VAF of the clone.

##### Insertion and deletion calling

For WGS data, small insertions and deletions were called using cgpPindel (*11*). The same matched normal samples were used for each donor, equivalent to the substitution calling with CaVEMan. In addition, several post-processing steps were carried out including (1) filtering of variants which did not pass the simple repeat filter (F017), (2) filtering of variants with a REP score > 3 and (3) filtering of variants with a reported VAF of 0.

##### Phylogenetic tree reconstruction

Phylogenies of single cell-derived AT2 organoids were generated as previously described and now implemented in the readily available R package Sequoia (https://github.com/TimCoorens/Sequoia) (*12*). In principle, the filtered substitutions from clonal AT2 organoids were used as input for MPBoot, creating a maximum parsimony algorithm (13). MPBoot was run with default parameters (command line: mpboot -s <ALIGNMENT file> -bb 1000, in which ‘bb’ denotes 1,000 bootstrap iterations) on concatenated nucleotide sequences of variant sites. The VAF was leveraged to represent the state of the mutations, where ‘1’ (present) was chosen for VAF ≥ 0.3, ‘0’ (absent) for VAF ≤ 0.1 and ‘?’ (unknown) for 0.1 < VAF < 0.3 to allow for uncertainty. An artificial nucleotide sequence represented by the reference genomes bases at variant sites was included to mirror the ancestral state at the zygote. The majority rule tree was utilised for mutation mapping relying on the calculation of the maximum likelihood for a mutation belonging to a branch implemented in the ‘treemut’ R package (https://github.com/NickWilliamsSanger/treemut).

##### Extraction of single-base pair substitution signatures

To identify mutational signatures for SNVs, the hierarchical Dirichlet process (https://github.com/nicolaroberts/hdp) on the 96 trinucleotide counts of all microdissected samples as well as to mutations assigned to each branch of the phylogenetic tree, was implemented. The HDP was run with individual patients as the hierarchy, in 20 independent chains, for 40,000 iterations and with a burn-in of 20,000. The identified components were compared to existing signatures and components with ≥ 0.90 cosine similarity were considered identical. The remaining signatures were deconvoluted using an expectation-maximisation algorithm, generally being explained by combinations of known signatures. In total, 9 signatures were evaluated which were then fitted back to the original mutation calls leveraging sigfit (https://github.com/kgori/sigfit).

##### Extraction of DBS and ID signatures

Indels detected in each sample were utilised to infer indel signatures using the MutationalPatterns package (*14*) in R. MutationalPatterns relies on non-negative matrix factorization to determine abundant signatures. All ID signatures detected previously (ID-1 - ID-18 (*15*)), were utilised as input for signature discovery. In total, 15 signatures were identified across all samples, although some were only abundant due to low indel burden samples.

##### Copy number calling from WGS

In order to identify copy number alterations (CNAs) in WGS, the Allele-Specific Copy number Analysis of Tumours (ASCAT) algorithm was (*16*) implemented within the ascatNGS package (https://github.com/Crick-CancerGenomics/ascat) (*17*). In brief, ASCAT was run with default parameters using a segmentation penalty of 100. Following this, ascatPCA, a custom-made filtering algorithm, was used. This algorithm reduces the number of false-positive calls that can arise when analysing genome sequences from normal tissue (https://github.com/hj6-sanger/ascatPCA). In principle, ascatPCA extracts a noise profile by aggregating the LogR ratio from a panel of normal, unrelated samples. Following this, the evaluated noise signature is subtracted from the signal observed in the sample being analysed, using principal component analysis.

##### Structural variant calling from WGS

Structural variation (SVs) were evaluated using GRIDDS (v. 2.9.4) with default settings (*18*). Based on the initial results, only SVs > 1kb in size and a quality score QUAL ≥ 250 were included. Furthermore, to increase the confidence in SVs < 30kb, an increased quality threshold of QUAL ≥ 300 was applied. SVs which had assemblies from both sides of the breakpoint were only considered if they were supported by at least four discordant and two split reads. Additionally, if imprecise breakends were detected, e.g. the distance between the start and end positions was smaller than 10bp), the respective SVs were filtered out. Moreover, a standard deviation filter based on the alignment positions of either SVs ends was implemented (discordant read pairs < 5) and removed potential germline variants and artefacts by using an in-house panel of 350 normal samples, where SVs which were found in more than 3 different samples in the panel of normals, were removed.

##### Selection analysis across WGS data

To systematically identify genes under positive selection in our and the basal cell dataset, dndscv in R was used (*19*). Initially, the global dN/dS ratios of all genes which were found to be mutated in our dataset was investigated. Genes with q-value<0.05 or p-value<0.001 as well as highly recurrently mutated genes (≥ 8 unique mutations) were considered as driver genes. In addition, genes previously reported in lung cancer or normal bronchial epithelium were leveraged for subsequent analyses (n = 51 genes) (*9, 20–22*). For the tissue specific analysis, SNVs were split according to tissue of origin and leveraged as input for dndscv. Gene-level dN/dS ratios were utilised to assess tissue specific selection, reporting point estimates if the p-value of the mutation type of interest was smaller than 0.05.

##### Variant calling from tNanoSeq data

The variant calling pipeline for NanoSeq has been extensively documented in the respective publications (*23, 24*). In contrast to conventional NanoSeq, which requires a matched normal to filter out germline SNPs, the variant calling for tNanoSeq is implemented without a matched normal sample, due to the high per base duplex coverage. Hence, the deduplicated bam file from the same samples was used as matched normal. For this purpose, deduplicated bam files were created by identifying and randomly sampling one read from each read bundle (RB; reads with the same barcode and mapping to the same position in the genome). Following this, SNPs were identified as mutations with a VAF ≥ 0.3 in comparison to the GRCh37 reference genome. A variant was considered a true positive mutation, if it fulfilled several criteria as listed in the following: (1) each RB (group of PCR duplicates) contains a minimum of two reads from each of the two original DNA strands; (2) the consensus base quality scores should be at least 60, ensuring strong support for a given base call; (3) the minimum difference between the primary (AS) and secondary alignment score (XS) should be higher than 50 to keep only read pairs with unambiguous mapping, essentially removing mapping artefacts; (4) the average number of mismatches in a group of reads should not be > 2, either in the deduplicated bam or samples itself; (5) the maximum number of 5’ clips needs to be 0; (6) the minimum number of improper read pairs needs to be 0 to avoid unreliable mappings; (7) base calls in read ends, referring to the last 8 bp from the 5’ or 3’ ends, are discarded due to unreliability; (8) for SNV calling, reads in the RB are not allowed to contain indels; (9) the minimum number of bulk reads per strand at a given site was required to be > 15x; (10) for a given mutation, the respective base will not be seen at a frequency > 0.01 in the matched normal; (11) a site should not overlap a common SNP and noise mask. The common SNP mask included a total of 27,205,965bp based on SNPs with an allele frequency (AF) > 0.001 in the 1000 Genomes project data. The noise mask was developed based on gnomAD indel calls with an AF > 0.01 and SNP calls with AF > 0.001 respectively. To create the mask, mismatches rates were calculated for every position across a panel of 448 in-house standard whole-genome sequences samples, resulting in a mask size of 22,474,160 bp in total.

##### Selection analysis across tNanoSeq data

The selection analysis for tNanoSeq data was implemented for AT2-enriched, AT2-depleted and all samples, respectively. Again, the dndscv package was utilised, albeit with additional covariates included. Dndscv on the targeted panel of genes was implemented, incorporating the average coverage per gene as well as a minimum coverage threshold of 50 dx. Moreover, a specific, tNanoSeq refCDS object was used, given that NanoSeq relies on filtering of common SNPs as mentioned above. In brief, common SNPs tend to be synonymous rather than non-synonymous, potentially inflating the dN/dS estimates for the nuclear genome to show positive selection. However, this artificial signal can be removed by masking the respective SNPs sites used for filtering. In addition, the refCDS object was updated by choosing a canonical isoform per gene using Appris (*25*) and variants of GENCODE and Ensembl gene annotations (*26*). Global dN/dS ratios of all genes were analysed in equivalence to WGS. Lastly, new implementations within the dndscv package were leveraged to investigate site- and codon-level selection in our data (*24*). Provided a list of common hotspots observed in TCGA and MSK cohorts, restricted hypothesis testing was conducted, specifically assessing selection on the provided sites. Significant sites were reported if they were in > 2 samples for sites and > 3 samples for codons and showed a q-value < 0.1.

##### Tumour-origin simulations under a multistage model of carcinogenesis

A substantial body of scientific literature has already explored the use of multistage models to contextualise cancer development. has established multistage models as a useful conceptual framework for understanding carcinogenesis. Building on this literature, the hypothesis that the difference in cancer risk between lung adenocarcinoma (LUAD) and lung squamous cell carcinoma (LUSC) arises from the distinct mutation burden distributions observed in alveolar and basal cells was examined using a bespoke simulation framework implemented in R (see Code availability). The framework was based on the classical Armitage–Doll multistage model of carcinogenesis, in which malignant transformation results from the sequential acquisition of *k* rate-limiting mutational events in a population of susceptible cells n₀ (*27*). Accordingly, and in simple terms, an individual’s cancer risk can be described as a function of the mutation rate *r*:

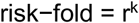

Although these models provide a deliberately simplified description of carcinogenesis and do not explicitly model spatial structure or detailed clonal ecology, they have proven conceptually valuable. They have previously been used to infer the number of rate-limiting steps in lung cancer, yielding estimates ranging from k = 1–6 for LUAD (*19*, *28*) and *k* = 3-11 for LUSC (*19*). Alexandrov and colleagues further demonstrated, using multistage simulations, that heterogeneity in mutation rates among normal lung epithelial cells can strongly shape cancer risk (29). Moreover, the relationship mentioned above has an important and often underappreciated consequence: when mutation rates vary across cells, tumour-initiating lineages are expected to be strongly biased toward cells with higher mutation rates, even if such cells constitute only a small fraction of the tissue. The magnitude of this bias depends critically on *k*. The simulation framework described below was designed to quantify this effect in both idealised settings and using empirically measured mutation rate distributions from normal human lung epithelium.

##### Conceptual tumour-origin simulations

To illustrate the statistical consequences of mutation rate heterogeneity under a multistage model independently of any specific biological system, a set of conceptual simulations was first performed using synthetic mutation rate distributions. A population of normal cells was generated consisting of two components: (i) a large population of low-mutation-rate cells sampled from a gamma distribution, and (ii) a much smaller subpopulation of hypermutator cells sampled from a second gamma distribution with a higher mean. This produced a bimodal distribution in which hypermutator cells represented approximately 1% of the population but contributed disproportionately to the upper tail of the mutation rate distribution. Given this population of cells, tumour-originating lineages were simulated by sampling cells with probability proportional to rᵏ, where *r* denotes the driver mutation rate of each cell and *k* is the assumed number of rate-limiting events. This sampling scheme directly implements the Armitage–Doll prediction that the probability of transformation scales as a power of the mutation rate. For each value of *k*, 5,000 tumour-originating cells were sampled, and the resulting distribution of mutation rates among tumour-initiating cells was compared to the underlying distribution in normal cells. This procedure was repeated for multiple values of *k* (e.g., k = 3, 5, 7) to illustrate how increasing the number of required driver events progressively amplifies the bias toward high-mutation-rate cells. For each condition, the full density distribution of driver mutation rates and the mean tumour-origin mutation rate were computed and visualised. These simulations serve purely as a conceptual demonstration of how rare high-mutation-rate cells can dominate tumour initiation under a multistage process, even when they represent only a small fraction of the tissue.

##### Empirical tumour-origin simulations using alveolar and basal epithelial data

The same framework was then applied to empirically measured mutation rate distributions from normal human lung epithelium. Per-cell driver mutation rates were estimated separately for alveolar (AT2) and basal epithelial cells and stratified by smoking status (never-smoker, ex-smoker, current-smoker). For this, we used the global dN/dS estimates from our WGS data to estimate the proportion of lung cancer driver genes (n = 86) under positive selection. The w_all_ estimates for alveolar and basal cells indicated that 31% of non-synonymous mutations in these genes are drivers in AT2 cells, whereas 78% of mutations are drivers in basal cells. The driver mutation rate was then multiplied by the fraction of driver genes in the total number of coding genes in the genome and by a factor of 0.01, based on the assumption that 99% of mutations are non-coding. Finally, the resulting driver mutation burden per cell was divided by the age of each patient to generate a driver mutation rate per cell per year. This rate was subsequently utilised in the simulation framework. For each combination of histology and smoking status, this empirical distribution of per-cell driver mutation rates was treated as the underlying population of normal cells. Tumour-originating lineages were then simulated by sampling mutation rates from this distribution with probability proportional to rᵏ, exactly as in the conceptual simulations. For each condition, 5,000 tumour-originating mutation rates were sampled. This yields the expected distribution of mutation rates among tumour-initiating cells under a multistage model, given the observed heterogeneity in normal tissue. The resulting tumour-origin distributions were compared directly to the underlying normal-cell distributions to visualise the magnitude of the selection bias imposed by the multistage process. To assess the sensitivity of this effect to the assumed number of rate-limiting steps, the procedure was repeated for multiple values of *k* spanning the plausible biological range for lung cancer. For each value of *k*, the mean tumour-origin mutation rate and the full density distribution were computed and visualised.

##### Repopulation and lifetime excess-risk dynamics in ex-smokers

To investigate how epithelial repopulation after smoking cessation could modify cancer risk trajectories over time, a separate set of simulations was performed to model lifetime excess risk under changing mutation rate environments. In these simulations, cancer risk was treated as a cumulative hazard that accrues over time. For never-smokers, the per-year driver mutation rate was assumed to be constant throughout life. For ex-smokers, the per-year driver mutation rate was assumed to be elevated from the age of smoking initiation until the age of cessation. After cessation, the mutation rate in ex-smokers was modelled as gradually converging toward the never-smoker rate via a repopulation process, implemented as an exponential relaxation toward the never-smoker mutation rate with a specified annual replacement fraction. At each age, the instantaneous hazard was assumed to scale as rᵏ, and cumulative risk was computed as the time integral of this hazard. Excess risk in ex-smokers was defined as the difference between the cumulative risk in ex-smokers and the cumulative risk in never-smokers. This excess risk was normalised to 100% at the age of smoking cessation, allowing direct visualisation of how risk accumulates during smoking and how it subsequently stabilises or declines depending on the rate of repopulation. This formulation correctly captures three qualitative features observed in epidemiological data: (i) excess risk increases approximately monotonically during the smoking period, (ii) excess risk plateaus or declines after cessation depending on repopulation dynamics, and (iii) the speed of post-cessation risk attenuation depends on the assumed rate of epithelial replacement by lower-risk clones.

#### Statistical analysis

Statistical analyses for experimental data were conducted using Graphpad Prism (v10.0.2) or R. Data are presented as mean ± standard error of the mean. p-values were calculated using a suitable statistical test as specified in the figure legends.

#### Data availability

Sequencing data will be made available upon publication of the manuscript on ENA (WGS: EGAD00001015821; tNanoSeq: EGAD00001015819). Curated lists of variant calls are provided on Zenodo (10.5281/zenodo.17828288).

#### Code availability

Computational methods provide a summary of the procedures implemented in various custom-made R, python and bash scripts. These scripts contain the commands run for the analyses highlighted in this publication. The code will be publicly available on Github upon publication (https://github.com/mjprzybilla/Alveoli_Manuscript).

### Supplementary Figures

**fig. S1.**
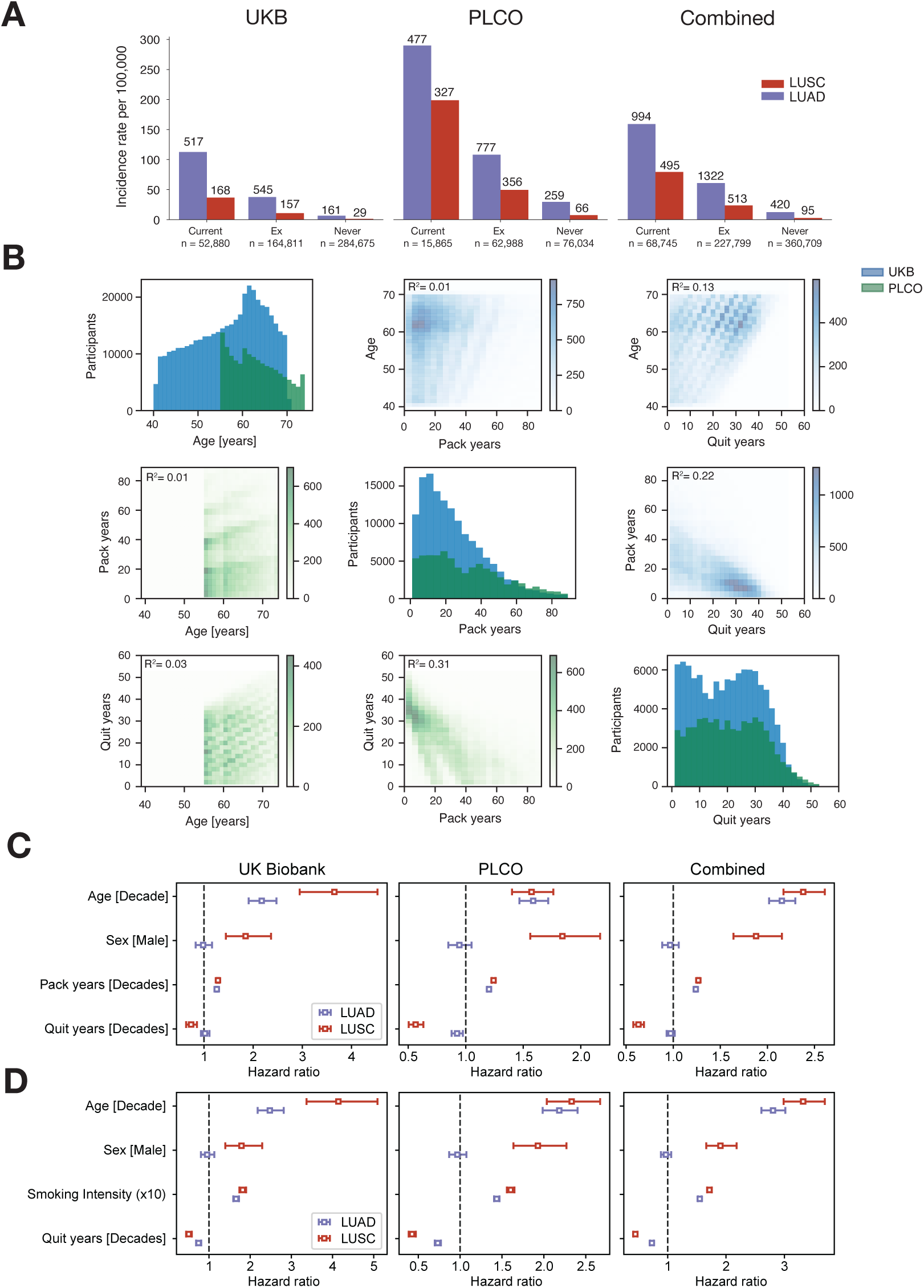
Extended Cox proportional hazard regression analysis in UKB, PLCO and combined datasets. **(A)**, Yearly rates and total number of LUSC and LUAD cases in UKB, PLCO and combined across smoking categories. (**B**), Summary of continuous regressors in Cox proportional hazards model (**C**), Hazard ratios associated with unit increases in each regressor, shown for each dataset. (**D**), Hazard ratios as in (**C**), for a model in which pack years are substituted for smoking intensity to reduce the correlation between regressors. Comparable results are seen.

**fig. S2.**
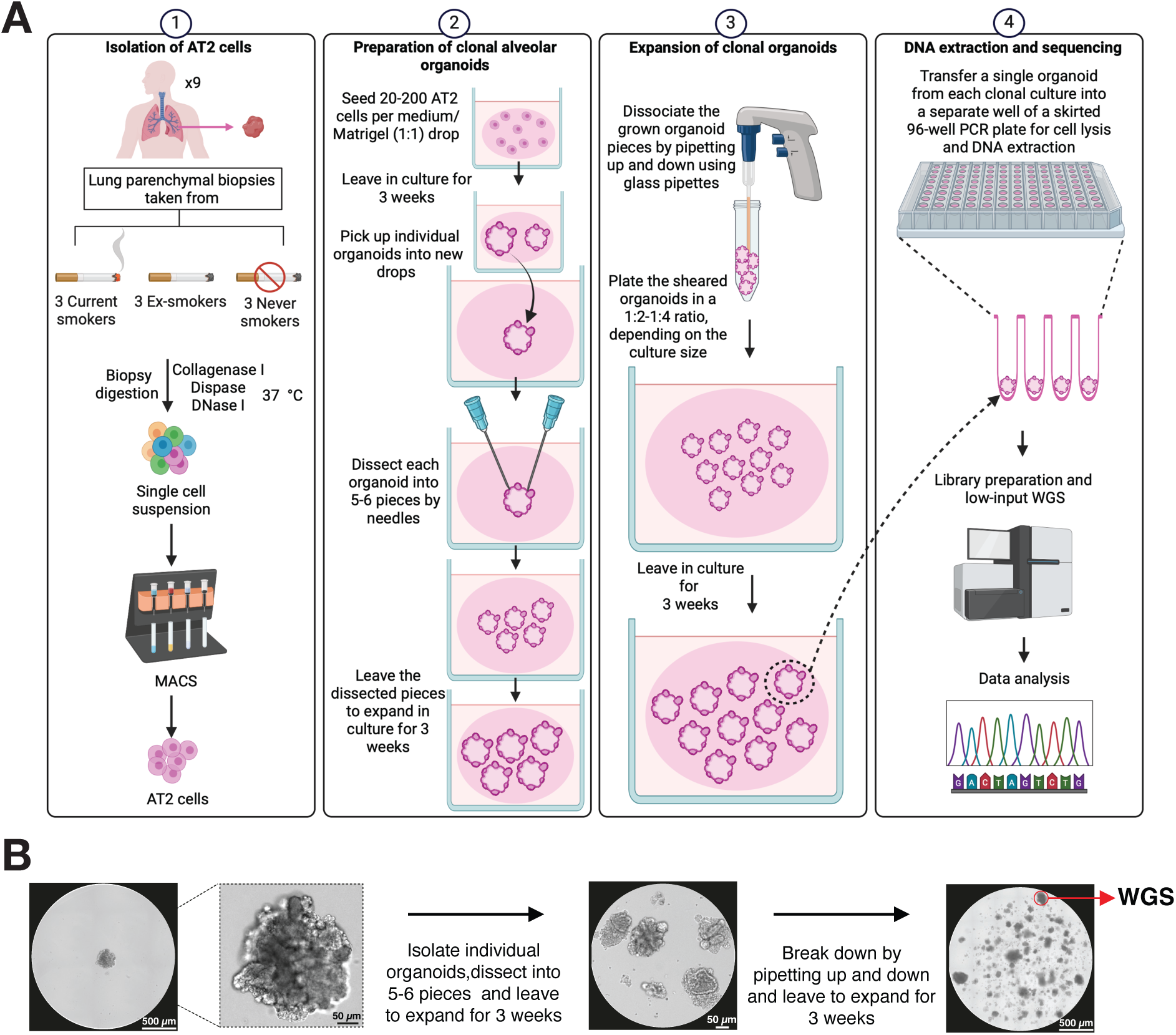
Experimental workflow for the creation of clonal alveolar organoids. (**A**) Schematic of the experimental process underlying the generation of clonal AT2 organoids. (**B**) Representative images of an established clonal organoid culture. The left image displays a drop containing a single grown organoid. To increase the chance of clonality, each organoid was passaged twice. In the first passage, the organoid was dissected into 5-6 pieces and left to grow in culture for 3 weeks (middle image). In the second passage, the grown organoid pieces were broken down by pipetting up and down and left to grow for 3 weeks to establish an organoid culture from which a single organoid was picked and utilized for whole-genome sequencing (right image).

**fig. S3.**
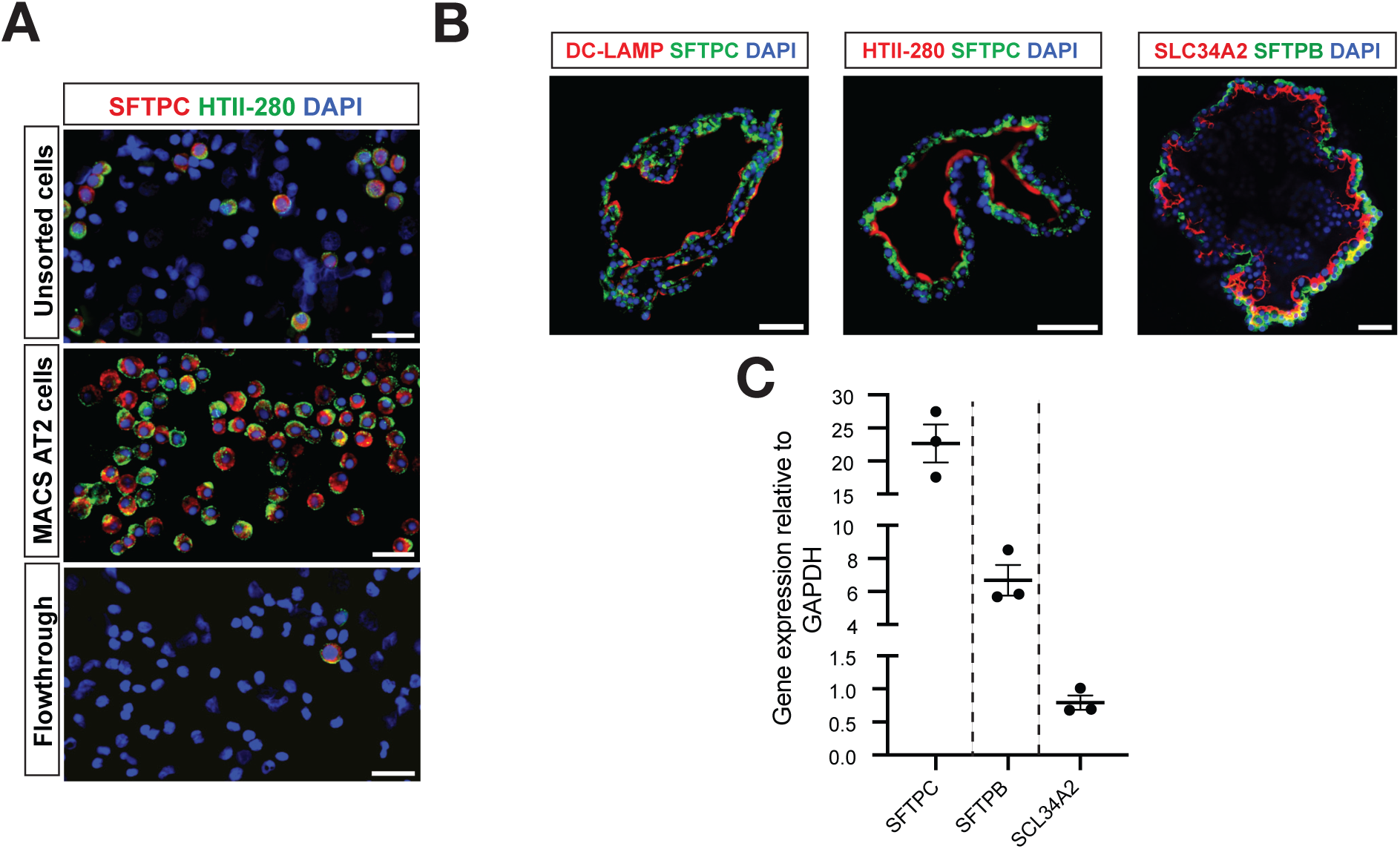
Experimental characterization of alveolar organoids. (**A**) Immunofluorescence images of cytospin preparations of the unsorted whole cell suspension (top) as well as the MACS-sorted AT2 cells (middle) and flowthrough (bottom). Sorted cells display expression of SFTPC (red) and HTII-280 (green), both of which are specific markers of AT2 cells. Scale bar = 20 µm. (**B**) Immunofluorescence images of alveolar organoids demonstrating expression of AT2 cell markers including SFTPC, DC-LAMP, HTII-280, SFTPB and SLC34A2. Scale bar = 50 µm. (**C**) Quantitative RT-PCR analysis for the expression of SFTPC, SFTPB and SLC34A2 genes in alveolar organoids, the chart displays the levels of gene expression relative to GAPDH. Relative gene expression was calculated using the 2^−ΔCT^ method. Depicted values represent the mean relative gene expression of each patient and error bars represent the standard error of the mean (n = 3 patients).

**fig. S4.**
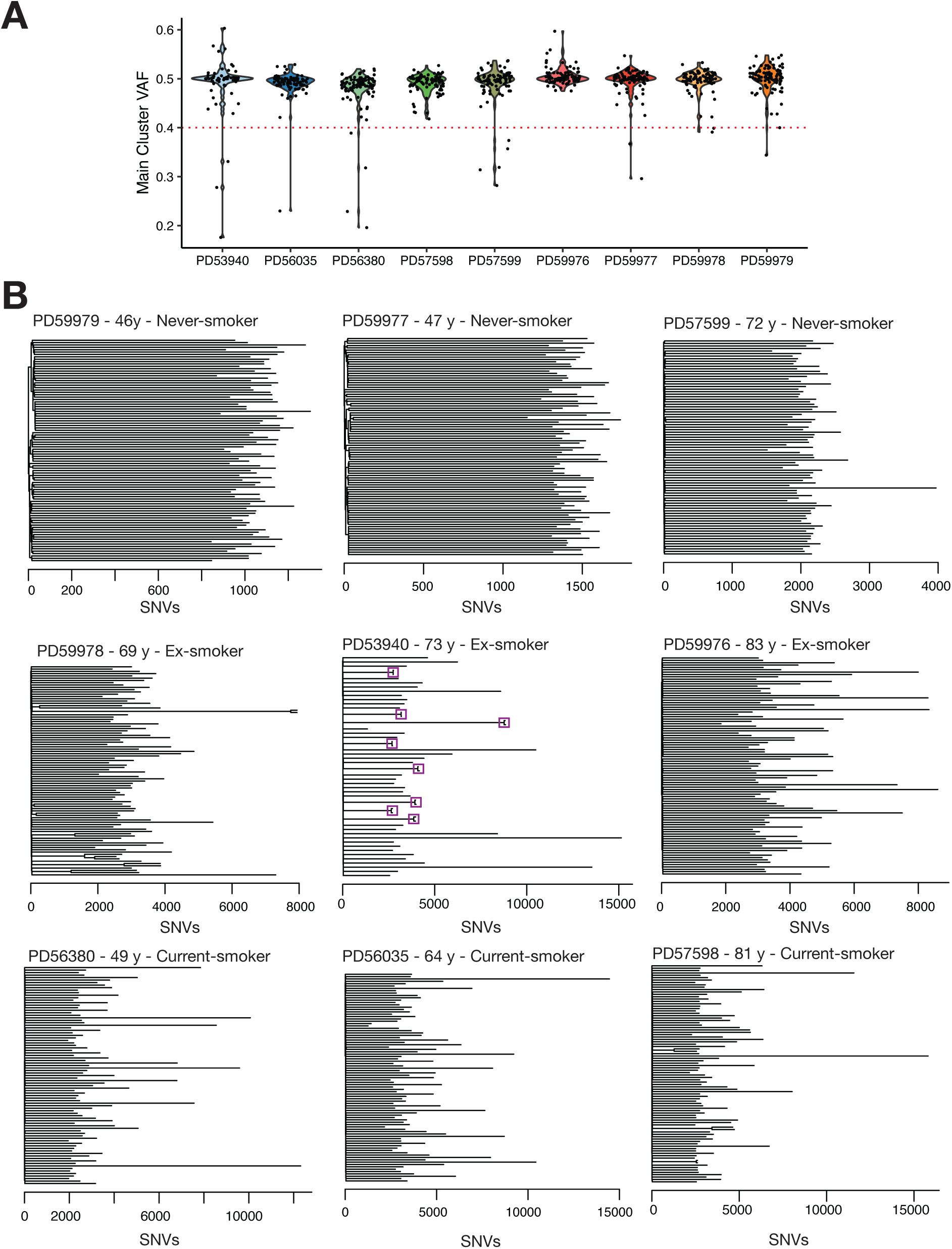
Clonality and phylogenetic relationships of alveolar organoids. (**A**) Violinplot depicting the results from the clonality estimation for each organoid across all patients. The variant allele fraction (VAF) of the main cluster inferred by the methodology is shown. The red dotted line represents the threshold for exclusion of organoids which are not monoclonal. (**B**) Phylogenetic trees showing the relatedness between clonal alveolar organoids across patients. Purple boxes indicate single cell-derived organoids from the same clonal expansion.

**fig. S5.**
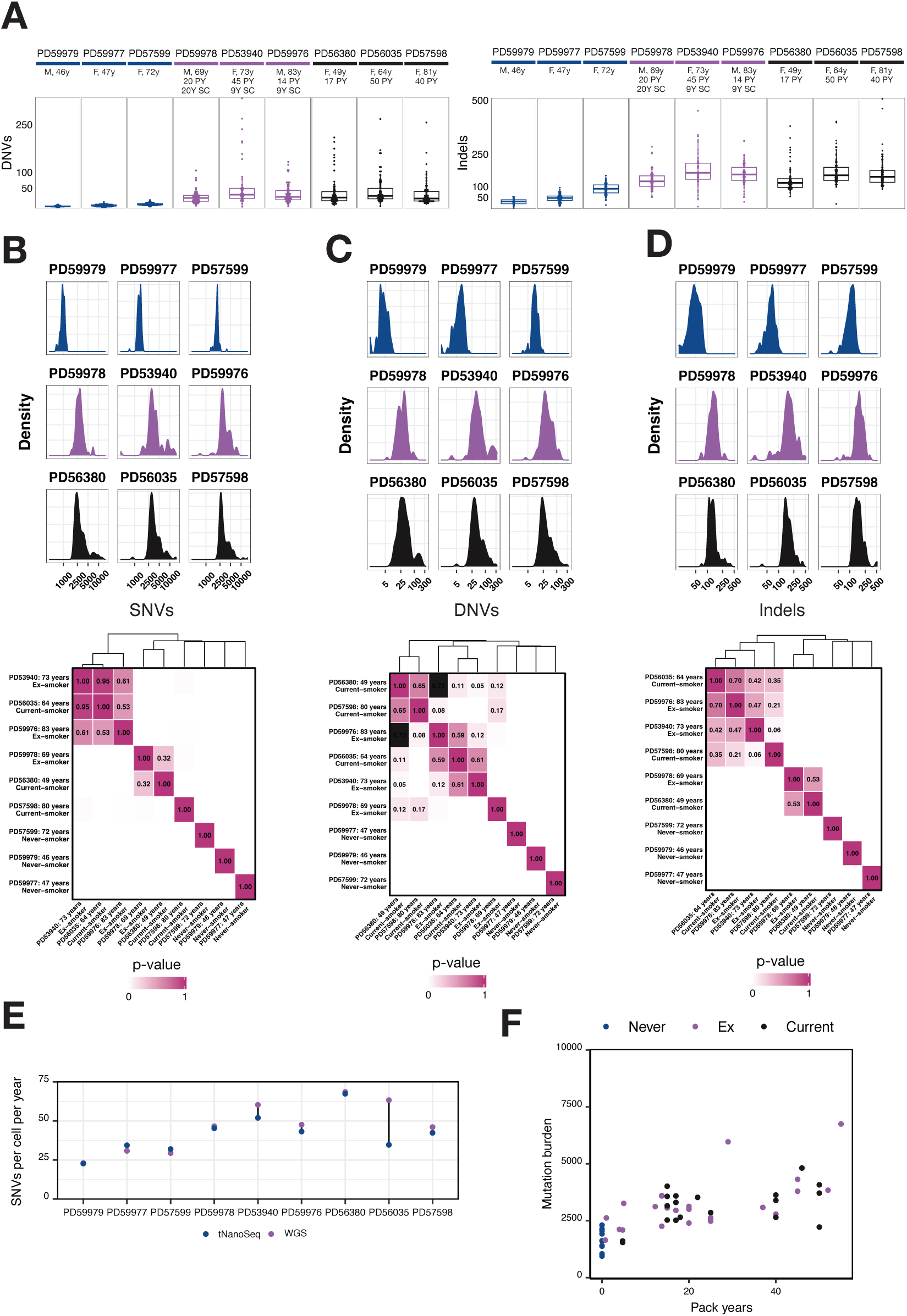
Analysis of genomic alterations in alveolar organoids. (**A**) Box plot depicting the total number of dinucleotide variants (DNVs, left) and insertions and deletions (indels) per AT2 cell (organoid) split according to the respective patient. Patients are ordered according to smoking status and age. Sex, age and smoking cessation as well as pack year history is annotated if applicable. (**B-D**), (top) Density distribution showing the burden of SNVs for each patient on a log transformed scale as well as (bottom) a heatmap depicting the results of a pairwise Kolmogorov-Smirnov (KS) test between the SBS burden distribution of each patient represented. A p-value > 0.05 demonstrates that there is insufficient evidence to conclude that the compared distributions differ. (**E**) Comparison of the number of SNVs per year per cell from targeted nanorate sequencing (tNanoSeq) and whole-genome sequencing (WGS) for the nine patients with alveolar organoids. (**F**) Scatterplot highlighting mutation burden compared to pack year history for all patients which underwent tNanoSeq or WGS. Smoking status is annotated for each point. PY: pack-years; SC: Smoking cessation.

**fig. S6.**
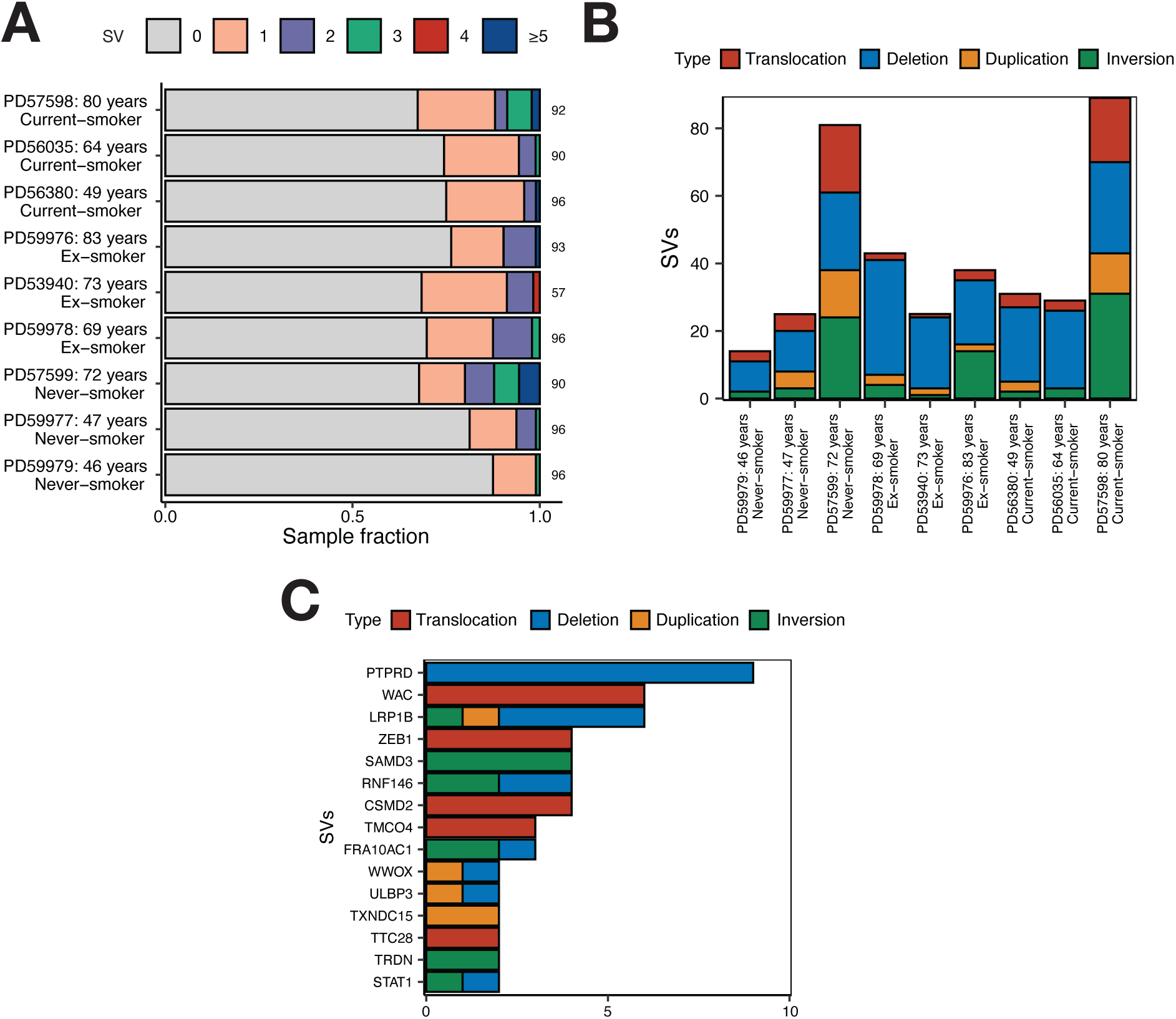
Structural variation across single cell-derived AT2 organoids. (**A**), Bar plot showing the proportion of samples affected by structural variation (SVs) per patient. For visualisation purposes, the number of alterations was capped at 5. (**B**), Number and type of SVs detected per patient. (**C**), Genes affected by deletions in single cell-derived AT2 organoids. Bar plot highlights the top 15 genes affected.

**fig. S7.**
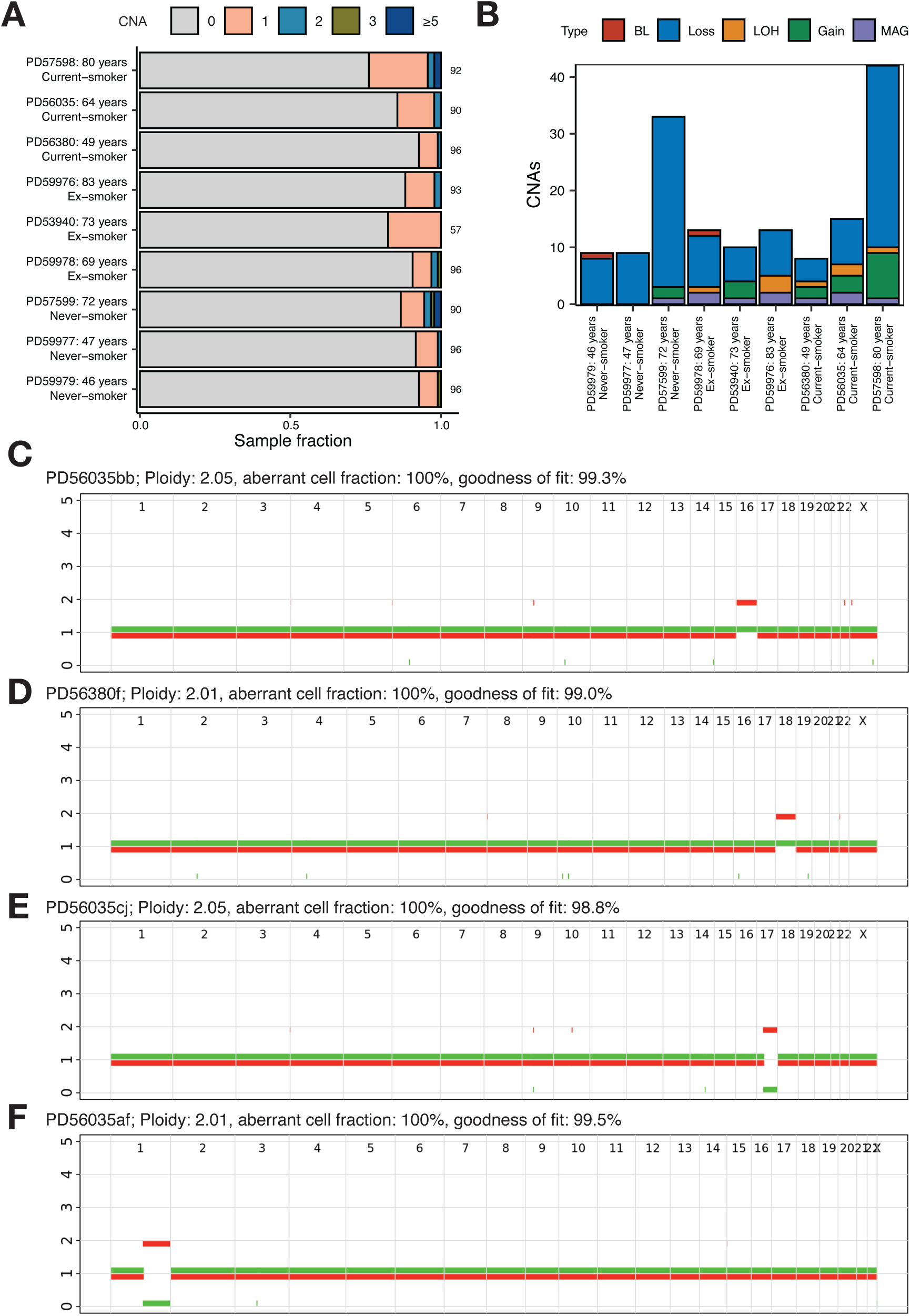
Copy number alterations across single cell-derived AT2 organoids. (**A**) Bar plot depicting the proportion of samples affected by copy number alterations (CNAs) per patient. For visualisation purposes, the number of alterations was capped at 5. (**B**) Number and type of CNAs detected per patient. BL = Biallelic loss, LOH = Loss of heterozygosity, MAG = Multi-allelic gain. (**C-F**) ASCAT plots presenting examples of CNAs observed in single cell-derived AT2 organoids.

**fig. S8.**
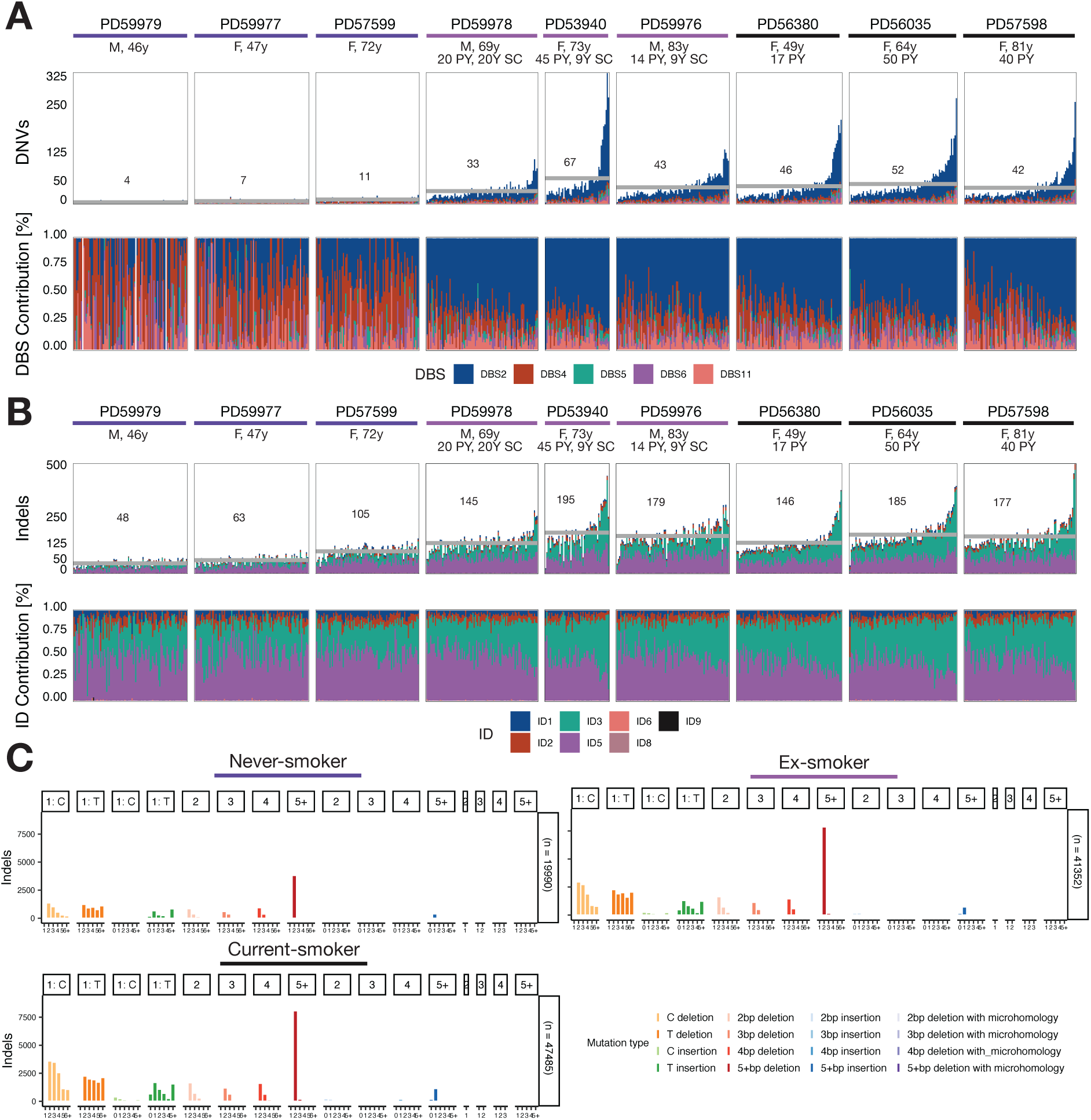
DBS and ID signatures in single cell-derived AT2 organoids. (**A**) (top) Bar plot depicting the total number of dinucleotide variants (DNVs) per AT2 cell coloured by DBS signature contribution as well as the relative contribution of DBS signatures (bottom). Visualisation is split according to the respective patient, with cells being ordered by increasing SNV burden as shown in Fig. 2. Patients are ordered according to smoking status and age. The horizontal grey line indicates the mean burden calculated across all organoids of a patient. Gender, age and smoking cessation as well as pack year history is annotated if applicable. (**B**) Equivalent to A for indels and ID signatures. (**C**) Indel spectra for never-, ex- and current smokers aggregated across all patients. The number of total indels detected is highlighted on the right.

**fig. S9.**
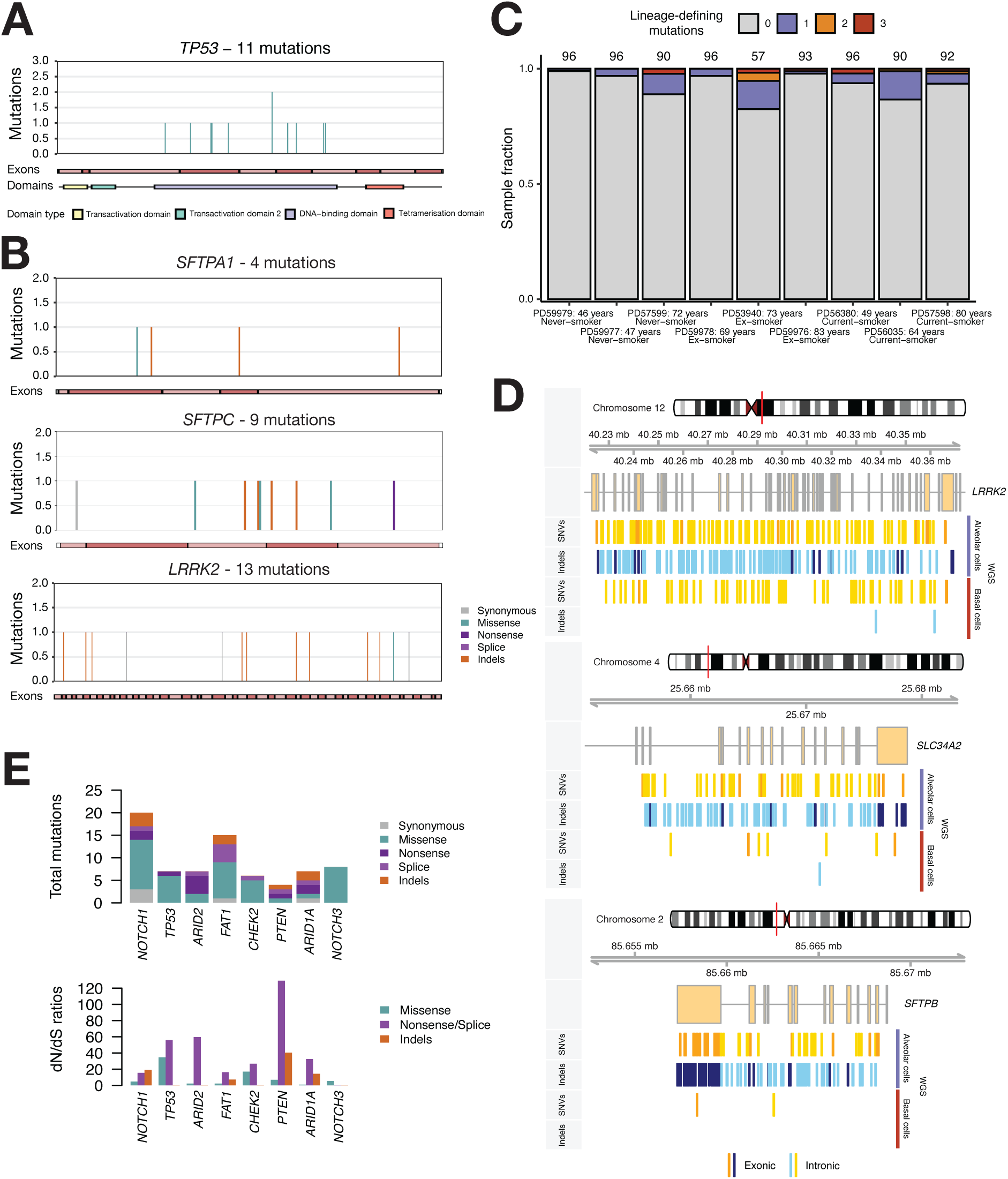
Selection analysis and lineage imprints across single cell-derived alveolar and basal cells. (**A**), Gene representation for *TP53* for mutations found across single cell-derived alveolar type 2 organoids. Number of mutations are annotated on the y-axis. Exon as well as protein annotation are shown if available for the respective transcript. (**B**), Equivalent representations as shown in (**A**) for *SFTPA1*, *SFTPBC* and *LRRK2.* (**C**), Barplot representing the fraction of cells per patient carrying a mutation in a surfactant protein (SFTP). The total number of samples per patient are denoted above each bar. (**D**), Genomic position of SNVs (top) and indels (bottom) for alveolar and basal cells detected in *LRRK2*, *SLC34A2* and *SFTPB*. Exonic and intronic indels are differentiated according to the respective colour. (**E**), Genes under positive selection in proximal airway data from Yoshida et al., 2020. (top), Number and consequence of mutations detected in WGS from single cell-derived basal cell colonies. (bottom) Observed-to-expected ratios for missense substitutions, truncating (nonsense and essential splice site) substitutions and indels. Genes shown are under significant positive selection (dndscv, q<0.1) when using lung cancer driver genes only.

**fig. S10.**
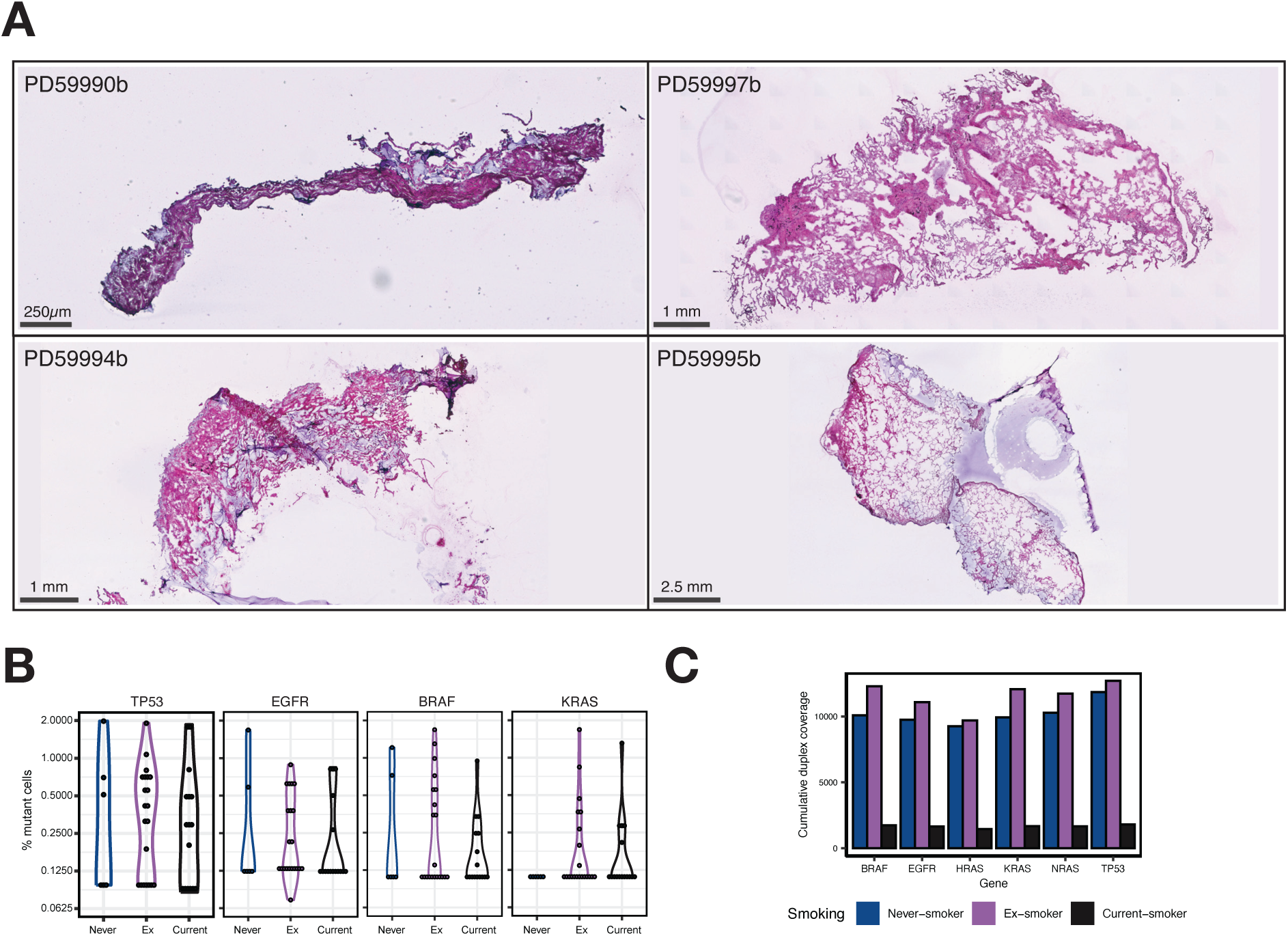
Genes under positive selection in all samples sequenced with tNanoSeq. (**A**), Examples for tissue histology of lung parenchyma samples used for tNanoSeq. (**B**), Fraction of mutated cells estimated from tNanoSeq for *TP53*, *EGFR*, *BRAF* and *KRAS*. Each individual dot highlights the fraction of cells affected by mutations in the respective gene per sample. Samples are split according to smoking status. All samples for each gene are plotted, even if no mutation was detected. In that case, the minimum cell fraction which could theoretically be detected was calculated using the mean duplex coverage per gene and the sample was set to the respective fraction of mutant cells. (**C**), Cumulative duplex coverage across all samples for *BRAF*, *EGFR*, *HRAS*, *KRAS*, *NRAS* and *TP53* across smoking categories.

**fig. S11.**
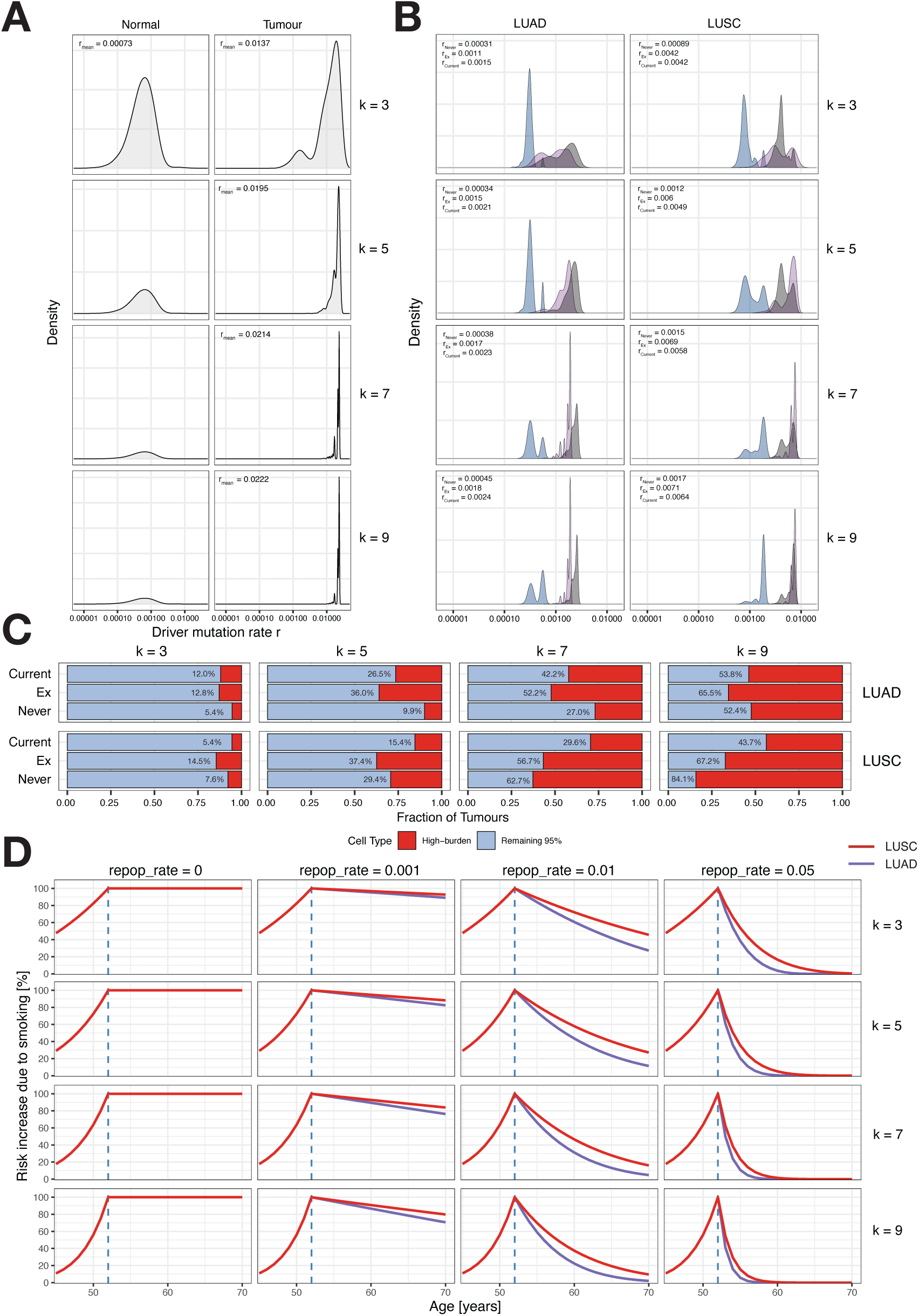
The influence of the number of events *k* on mutation burden and risk. (**A**), Simulated driver mutation rate distribution of normal cells (top) and for cancer (bottom) with *k* = 3, *k* = 5, *k* = 7 and *k* = 9 events. Note a small population of cells with high mutation burden in the top panel. (**B**) Simulated tumour origin mutation rate *r* from the empirical data for *k* = 3, *k* = 5, *k* = 7 and *k* = 9 events. (**C**) Bar plot highlighting the proportion of tumours developing from the 5% high mutation burden cells compared to 95% of the remaining cells in our empirical data. Varying number of events are shown for both LUAD and LUSC. Annotated percentages show the percentage of tumours resulting from hypermutator cells. (**D**) Simulated risk over time for LUAD and LUSC according to an Armitage-Doll model. Risk increase is simulated as excess risk over a never-smoker. Both *k* and the repopulation rate (*repop_rate*) are varied across panels.

### Supplementary Tables

**Table S1.** Extensive patient information and metadata.

**Table S2.** QC metrics and metadata for single cell-derived AT2 organoids from WGS.

**Table S3.** Copy number alterations and structural variants detected in single cell-derived AT2 organoids from WGS.

**Table S4.** QC metrics and metadata for alveoli samples from targeted NanoSeq.

**Table S5.** Primary antibodies.

| Antibody | Vendor | Cat. No. | Dilution |
| --- | --- | --- | --- |
| Mouse monoclonal anti-SFTPb | Thermo Fisher Scientific | MA1-204 | 1:400 |
| Rabbit polyclonal anti-SFTPb | Thermo Fisher Scientific | PA5-42000 | 1:400 |
| Rabbit polyclonal anti-SFTPC | Abcam | ab90716 | 1:200 |
| Rabbit polyclonal anti-SFTPC | Merck Millipore | Ab3786 | 1:400 |
| Mouse monoclonal anti-HTII-280 | Terrace Biotech | TB-27AHT2-280 | 1:250 |
| Rat monoclonal anti-DC-LAMP | Novus Biologicals | DDX0191P | 1:250 |
| Rabbit monoclonal anti-SCL34A2 | Abcam | ab228474 | 1:100 |
| Chicken polyclonal anti-KT5 | Biolegend | 905901 | 1:500 |
| Rabbit monoclonal anti-TP63 | Abcam | ab124762 | 1:400 |

**Table S6.** Secondary antibodies.

| Antibody | Vendor | Cat. No. | Dilution |
| --- | --- | --- | --- |
| Donkey anti-mouse<br>Alexa Fluor 555 | Thermo Fisher Scientific | A31570 | 1:1000 |
| Donkey anti-rabbit<br>Alexa Fluor 647 | Thermo Fisher Scientific | A31573 | 1:1000 |
| Donkey anti-rat<br>Alexa Fluor 488 | Thermo Fisher Scientific | A21208 | 1:1000 |
| Donkey anti-rabbit<br>Alexa Fluor 555 | Thermo Fisher Scientific | A31572 | 1:1000 |
| Goat anti-chicken<br>Alexa Fluor 647 | Thermo Fisher Scientific | A21449 | 1:1000 |
| Donkey anti-rabbit<br>Alexa Fluor 444 | Thermo Fisher Scientific | A21206 | 1:1000 |
| Goat anti-mouse IgM Alexa<br>Fluor 488 | Thermo Fisher Scientific | A10680 | 1:1000 |
| Donkey anti-mouse Alexa<br>Fluor 647 | Thermo Fisher Scientific | A31571 | 1:1000 |

